# Tissue Fluidity Mediates a Trade-off Between the Speed and Accuracy of Multicellular Patterning by Cell Sorting

**DOI:** 10.1101/2025.03.01.640992

**Authors:** Rikki M. Garner, Sean E. McGeary, Allon M. Klein, Sean G. Megason

## Abstract

The organization of cells into spatial patterns is a fundamental aspect of multicellularity. One major mechanism underlying tissue patterning is adhesion-based cell sorting, in which a heterogeneous mixture of cell types spontaneously separates into distinct domains based on differences in adhesion protein expression. Here, we identify tissue fluidity—the extent to which cells can move freely within a tissue—as a critical regulator of adhesion-based sorting. First, we describe a physically well-understood minimal tissue model that can integrate both tissue fluidity and adhesion-based sorting, and demonstrate that this model can quantitatively reproduce experimentally measured sorting dynamics in a fibroblast cell culture assay. We go on to show that altering tissue fluidity by any mechanism in the model leads to substantial changes in the rate or accuracy of sorting (or both). We further demonstrate that the balance between cell motility, which acts to fluidize the tissue, and homotypic cell–cell adhesion, which acts to solidify the tissue, sensitively tunes a fundamental trade-off between the rate and accuracy of sorting—such that sorting can only occur when motility and adhesion are tightly coupled. Intriguingly, best fits of the simulations to the experiments across a range of adhesion protein expression conditions suggest that cells may naturally scale their motility strength with their adhesion strength – thereby maintaining a permissive fluidity for sorting. Overall, our results indicate that tissue fluidity must be tightly regulated for sorting to occur, and that cells may have evolved a mechanism to naturally co-regulate their mechanical properties in order to sustain a patterning-competent fluidity.

**Statement of Significance:** Tissue fluidity, or the ability of cells to freely rearrange within a tissue, is a universal property of multicellular organisms that plays central roles in development, cancer, and wound healing. Here, we identify tissue fluidity as a critical regulator of a major mechanism of multicellular patterning – adhesion-based cell sorting. The results of our combined experimental-computational investigation suggest that tissues can readily tune their fluidity in order to freeze, catalyze, or erase multicellular patterns – carrying significant implications for our understanding of how patterns are formed in development, lost in diseases affecting tissue organization (e.g., cancer), regained through the processes of wound healing and regeneration, and can be engineered in the creation of synthetic organoid and embryoid systems.

**One sentence summary:** Biophysical modeling demonstrates how tissue fluidity is a key regulator of the rate and accuracy of adhesion-based sorting.

## Introduction

The precise spatial patterning of cells within tissues is essential to multicellular life. Given its broad importance, a general understanding of how multicellular patterns arise is foundational to nearly every aspect of tissue and organismal biology – including how patterns are formed in development, lost in diseases affecting tissue organization (e.g., cancer), regained through the processes of wound healing and regeneration, and engineered in the creation of synthetic organoid and embryoid systems. Classical views of cell type patterning (e.g., Lewis Wolpert’s French flag(*1*)) imagined cells as stationary objects whose fates are determined by an external signaling gradient. However, modern time-lapse imaging approaches have revealed that cells often undergo a significant amount of both directed and random migration within developing tissues(*2–5*), and that morphogen signaling and gene expression themselves can be noisy(*2*, *6*– *8*). Moreover, recent successes in stem cell and explant models of development have shown that minimal cell systems can self-organize to recapitulate many aspects of embryogenesis—including germ layer patterning(*9*, *10*), gastrulation(*11*), neural tube morphogenesis(*12*), somitogenesis(*13*), and head formation(*9*)—in the absence of the endogenous embryonic context. Together, these observations demonstrate a critical need for new conceptual frameworks to understand how tissue patterns can arise from cellular self-organization(*10*, *14*).

One major mechanism of cellular self-organization with important roles in development is cell sorting. Cell sorting is the process by which a heterogeneous mixture of different cell types spontaneously separates out into distinct domains, typically due to differences in physiological properties between the cell types. Based on our work and others, it is clear that much of embryonic patterning relies on cell sorting(*15*) based on differences in signaling state(*8*) and cell biomechanical properties such as adhesion protein expression(*8*, *16–22*), cortical tension(*23*), or cell shape dynamics(*24*).

But while it is clear that cells *can* sort based on physiological properties, we lack a detailed understanding of how cells move through the tissue, find cells of the same cell type, and then ultimately stop moving to lock in the sorted pattern. In other words, we have much to learn about how biological regulation of cell motility and related biomechanical properties controls cell sorting. Here, we investigate the role of tissue fluidity (the ability of cells to rearrange within a tissue) in adhesion-based sorting. A deep body of literature on tissue rheology(*23*, *25–34*) has revealed that a diverse range of tissues in a variety of contexts have been shown to transition between a fluid-like state, where cells can rearrange easily, and a more solid-like state, where cells are constrained in place(*25–28*, *30*, *35*, *36*). During development, transitions in tissue fluidity drive crucial morphogenetic events such as body axis elongation(*35*, *36*). Changes in tissue fluidity have also been implicated in wound healing(*37*) and diseases including cancer(*23–26*) and asthma(*38*). A predictive, mechanistic understanding of the relationship between tissue fluidity and cell sorting is thus critical to our understanding of these central biological processes.

Physics has much to offer biology in this regard. Biophysical models of cell sorting have been developed extensively over the past decades(*29*, *39–46*), identifying important roles for many physical tissue properties associated with tissue fluidity(*47*), including cell motility(*39*, *40*, *45*, *46*). Given the recent surge of interest in tissue fluidity and cell sorting across diverse physiological and synthetic contexts, it is essential to revisit these ideas in direct practical comparisons with experimental measurements of cell sorting – and to evaluate the constraints for cell sorting on biologically-relevant length- and time-scales.

In this work, we apply an established statistical physics formalism to demonstrate the basic physical principles driving the interplay of tissue fluidity and cell sorting. After demonstrating that this model can quantitatively recapitulate the sorting patterns and dynamics observed in experiments, we show that (1) tissue fluidity is a critical regulator of adhesion-based sorting, (2) the balance of two major determinants of tissue fluidity—cell motility and cell–cell adhesion— finely tunes a fundamental trade-off between the rate and accuracy of sorting, with an intermediate fluidity being optimal for sorting, and (3) cells may intrinsically scale their strength of motility with their adhesion strength in order to robustly maintain a permissive fluidity for sorting.

## Results

### A minimal biophysical model of tissue fluidity and adhesion-based sorting quantitatively predicts sorting dynamics in an experimental cell sorting assay

To interrogate the relationship between tissue fluidity and cell sorting, we implemented a minimal biophysical model of a tissue (**Fig. 1**, see ***Methods***). The model takes the form of a Kawasaki Ising grid model(*48–51*), a well-established approach in statistical physics for modeling biological tissues(*39*, *52*). There exist many diverse definitions and uses of the term tissue fluidity(*25*, *35*, *37*, *47*, *53–57*). Tissues are collective cellular systems and thus inherently multi-scale, and so tissue fluidity can, and has been, defined at the tissue level (e.g., viscosity and the resistance of the tissue to flow(*25–29*, *35*, *55*, *56*, *58*, *59*)), the meso-scale (e.g., rigidity percolation(*25*), jamming/unjamming transitions(*38*, *54*, *60*)), and the cell scale (e.g., neighbor exchange(*30*, *35*, *37*, *57*), cell motility(*29*, *35*, *36*, *58*)). Here, for the context of understanding cell sorting, we define tissue fluidity as the ability of cells to rearrange within the tissue (*35*, *37*). We quantify tissue fluidity as the rate of cell rearrangement, such that cells are stationary in a solid tissue, and cell rearrangement increases with increasing tissue fluidity (**Fig. 1a-c**). Tissue fluidity in the model is set by a competition between cell–cell adhesion and cell motility (**Fig. 1b**). Cell–cell adhesion resists cell rearrangement, so tissue fluidity decreases with increasing cell–cell adhesion strength. On the other hand, the propulsive force of cell motility provides the energy for the cell to overcome the adhesion forces and exchange places with their neighbor, so tissue fluidity increases with the strength of motility (**Fig. 1c**).

**Figure 1.**
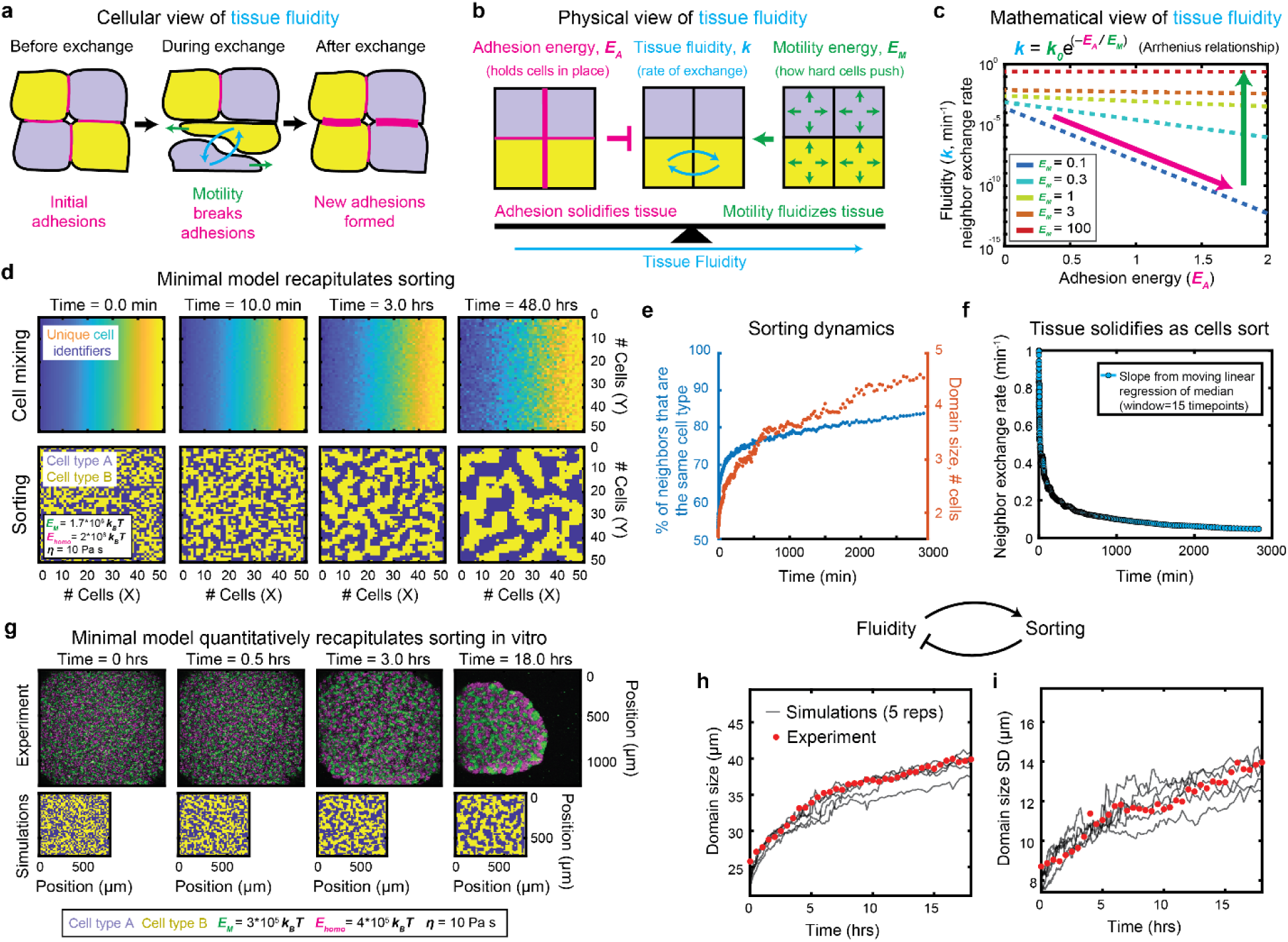
A minimal dynamic model to study the interplay between tissue fluidity and adhesion-based sorting. (**a**) Schematic of the model at the cellular level. (**b**) Schematic of the model’s physical interpretation of tissue fluidity. (**c**) Schematic of the model’s mathematical interpretation of tissue fluidity. (**d**) Time-lapse montage of an example simulation initialized with two cell types randomly mixed in equal proportion, under the case of equal homotypic adhesion between cells of the same cell type and no heterotypic adhesion. *Top:* Color represents a unique identifier for each cell, determined by that cell’s initial position. The mixing of cells in space is indicated by the increasing rearrangement of colors over time. *Bottom:* Color represents each cell’s cell type, where sorting can be visualized by the formation of spatial domains of the same cell type. (**e**) For the simulation shown in (d), the degree of sorting is plotted as a function of time, using two different metrics of sorting. The left-hand y-axis represents the fraction of each cell’s four direct neighbors (i.e., sharing faces) that are the same cell type as that cell, averaged across all cells in the tissue. The right-hand axis represents the average width of domains of the same cell type in the tissue (see ***Methods***). (**f**) The tissue fluidity (i.e., the rate of neighbor exchange averaged over the tissue) over time (see ***Methods***). (**g**) Time-lapse montage of the experimental cell-sorting assay (*top*) and its associated best-fit simulation (*bottom*). *Top:* L929 cells co-expressing either *Cdh3* (P-cadherin) and *EGFP* (green) or *Cdh1* (E-cadherin) and *mRFP1* (magenta) were mixed in equal proportions and imaged by confocal time-lapse microscopy. Images represent maximum intensity projections. *Bottom:* Best-fit simulations displayed as a heat map of cell type. Best fit parameters are listed in the inset legend. (**h-i**) Mean (h) and standard deviation (i) of the same-cell-type domain size over time for the experiments (red dots) and for 5 replicate best-fit simulations (grey lines).

More specifically, we treat cell–cell adhesion as an energy barrier (***E_A_***) needed to be overcome for cells to swap places, and we treat cell motility as the energy (***E_M_***) that cells exert to overcome this adhesion energy barrier. Assuming cell motility in the tissue is random, we model cell motility as an effective thermal energy (arising from an effective temperature), as previously (*36*, *40*, *45*). The model is agnostic to the specific molecular or cellular mechanisms underlying the random cell motility, but could, for example, represent the generation of actin-rich or bleb-based protrusions (**Fig. 1c**) or dynamic fluctuations at cell–cell interfaces(*53*).

Borrowing from chemical kinetics, we assumed the simplest possible relationship between the rate of neighbor exchange (i.e., tissue fluidity, ***k***) and the adhesion and motility energies—a diffusion-limited Arrhenius relationship (***k*** = ***k_0_*** exp(–***E_A_*** / ***E_M_***), **Fig. 1c**, ***Methods***). The prefactor, ***k_0_***, represents the neighbor exchange rate in the absence of cell–cell adhesion (***E_A_***=0, ***k*** = ***k_0_***). As cells can crawl through tissues by many mechanisms that don’t require cell–cell adhesion (e.g., ECM-mediated motility, amoeboid migration, bleb-based motility), the energy exerted by cells to move (***E_M_***) is expected to drive cell rearrangements even in a tissue with no cell–cell adhesion (***E_A_***=0, ***k*** = ***k_0_***). Therefore, we chose ***k_0_*** to be proportional to ***E_M_***. More specifically, analogous to chemical kinetics, we chose ***k_0_*** to be a diffusion-limited neighbor exchange rate (***k_0_ = E_M_ / 3πηd^3^***), where ***η*** is the viscosity and ***d*** is the cell diameter. Therefore, just as with thermal energy in chemical kinetics, the motility energy appears in both the prefactor (***k_0_***), because it provides the energy for cells to move in the absence of adhesion, as well as the exponent, because it provides the energy to overcome cell–cell adhesion. Here, the effective viscosity, ***η***, experienced by the cell represents resistance to cell flow due to effects other than adhesion (e.g., viscoelastic cell deformation, friction between neighbors, extracellular matrix, extracellular fluid viscosity). Hence, increasing viscosity reduces tissue fluidity.

Cell–cell adhesion strength is dependent both on each individual cell’s adhesion protein expression profile as well as the immediate context of its neighbors. Therefore, in our model we define adhesion strength, and thus tissue fluidity, separately for each individual cell at each moment in time (taking into account neighbor information). This cell-level view of tissue fluidity can account for heterogeneity in the tissue(*29*)—an essential aspect of sorting—in contrast to commonly used bulk metrics for tissue fluidity (e.g., viscosity values derived from micropipette aspiration experiments on tissues). Indeed, similar cell-level modeling approaches have shown that cell–cell heterogeneity in growth and active fluctuations are advantageous for sorting(*29*). Our cell-level metric of fluidity can also be averaged across all cells within a tissue to arrive at a bulk measurement of tissue fluidity (**Fig. 1d-f**).

We next incorporated these elements into a stochastic simulation (see ***Methods***), where two different cell types with defined homotypic (same-cell-type) adhesion and cell motility are mixed. The model is equivalent to an Ising grid model with Kawasaki dynamics (*48–51*), implemented with a fixed discrete time step (see ***Methods***). We find that this model gives rise to both tissue fluidity (i.e., cell rearrangements over time, **Fig. 1d, top**) and cell sorting (**Fig. 1d, bottom**) – allowing us to interrogate the relationship between them (**Fig. 1e-f**). We quantified the degree of sorting as the fraction of each cell’s four direct neighbors (i.e., sharing faces) that were the same cell type as the cell of interest (**Fig. 1e, *left***), and as a more intuitive metric we also provide the average width of the same cell type domains (**Fig. 1e, *right***). The two complementary metrics scale monotonically with each other (**Fig. 1e**). We calculated the neighbor exchange rate as the number of neighbor exchanges per cell per unit time (**Fig. 1f**). By quantifying the degree of sorting and the average neighbor exchange rate over time (**Fig. 1 – Supp.** Fig. 1, see ***Methods***), we find that sorting initially progresses quickly while the tissue fluidity is high. However, as cells sort, the overall adhesion strength in the tissue increases, causing the tissue to solidify over time (**Fig. 1f**). The solidification of the tissue in turn coincides with a rapid slow-down in the sorting rate (**Fig. 1e**).

This general slowing down of phase ordering (here, sorting) and neighbor rearrangements over time are well-known features of the Kawasaki Ising model. However, the classic behavior of the model is such that the domain size grows with time as a power law of t^1/3^ in the long-time regime(*48*). Interestingly, we find that sorting for biologically-relevant timescales and parameter choices occurs within the low-time metastable glassy regime, where growth exhibits sub-t^1/3^ scaling with dynamics that are very sensitive to the motility energy (**Fig. 1 – Supp.** Fig. 2). These metastable glassy dynamics were previously observed in Kawasaki Ising models in the context of ferromagnets, which showed that phase separation dynamics are very temperature-sensitive in this regime (*61*). Glassy dynamics have also been reported in other models of biological tissues (*58*, *60*, *62*).

To determine the predictive power of our model, we turned to an experimental cell-sorting assay (see ***Methods***). For these experiments, L929 fibroblast cells, which do not endogenously express cadherins(*63*), were used to generate stable lines co-expressing either *Cdh1* (E-cadherin) and *mRFP1* or *Cdh3* (P-cadherin) and *EGFP. C*ells from either cell line were trypsinized, mixed in equal proportions, pelleted onto round-bottom 384-well plates, and imaged via time-lapse confocal microscopy. The cells in the experimental cell-sorting assay produced patterns which closely resembled those of the biophysical model (**Fig. 1g, top and Fig. 1d, bottom**). To quantify sorting, we measured the average (**Fig. 1h**) and standard deviation (**Fig. 1i**) of the same-cell-type domain widths in the experiments over time, using an identical analytical pipeline to that used for the simulations (**Fig. 1e**, **Fig. 1 – Supp.** Fig. 1). Note that we restrict the analysis of the experimental system to the first 20 hours of sorting. After this time, the aggregates start to break apart in a heterogeneous fashion and are no longer amenable to a sorting analysis – perhaps related to burst-like spreading and asymmetric domain expansion observed in other sorting tissues(*64*).

The form of the experimentally observed pattern can be used to infer the relative magnitudes of the homotypic and heterotypic adhesion energies(*17*). In particular, the finger-like pattern suggests that the homotypic adhesion energies are much larger than the heterotypic adhesion energies (***E_A,homo_*** > ***E_A,het_***). In contrast, if the heterotypic adhesion energy were to dominate or be comparable to the homotypic adhesion energies, one would have expected a checker-board like pattern or an inside-outside pattern (i.e., engulfment of one cell type by another). Additionally, the fact that the domain sizes of the two cell types being mixed were very similar to each other suggests that the homotypic adhesion energies of the two cell types were very similar (E_A,homo,11_ ∼ E_A,homo,22_). While our model can be used to explore sorting under any of these regimes, in this work we constrained further analysis to the regime most consistent with the experimental results (***E_A,homo,11_*** ∼ ***E_A,homo,22_*** > ***E_A,het_***). For fitting the model to the experimental data, we further made the simplifying assumption that both cell types being mixed had equal homotypic adhesion energy, and that heterotypic adhesion energy was zero. Therefore, in cases where two cell types with slightly different homotypic adhesion energies are mixed in experiments, the best fit model returns a single value representing an approximate average homotypic adhesion energy between the two cell types.

In order to determine the simulation parameters which best quantitively recapitulated the experimental data, we ran sorting simulations spanning a broad range of physiological parameters for the adhesion energy, motility energy, and viscosity (see ***Methods***). We then performed magnitude-weighted least absolute difference optimization to identify the parameter set which best matched the experimental sorting dynamics (**Fig. 1g-I**, see ***Methods*)**. The best-fit simulation recapitulated the experimentally-observed patterning in striking detail—reproducing not only the visual pattern (**Fig. 1g, Video 1**) but also the mean and variance in the domain size, as well as their dynamics over time (**Fig. 1h-i**).

The best-fit model allows us to estimate the relative values of the motility energy (***E_M_***), adhesion energy (***E_A_***), and viscosity (***η***) of the experimentally observed tissue. Because tissue fluidity in the model is determined by two major ratios—the diffusion limited neighbor exchange rate (***k_0_ = E_M_ / 3πηd^3^***) and the exponent (***E_M_ / E_A_***)—different parameter choices with identical ratios will yield an identical sorting behavior and so cannot be distinguished by the model. Therefore, given the best-fit ratios, we can predict two of the three parameters, after assuming a given value for the third. Assuming a viscosity of 10 Pa·s(*25*), the model predicts that the energy of motility is 3*10^5^ ***k_B_T***, comparable to the homotypic adhesion energy at a single cell–cell interface of 4*10^5^ ***k_B_T***. These estimates are well-within the experimentally-measured range of values in similar systems as well as back-of-the-envelope calculations (see ***Methods***). We note that extracting the individual values for ***E_M_*** and ***E_A_*** requires the model assumption that the adhesion-free neighbor exchange rate (i.e., the prefactor ***k_0_ = E_M_ / 3πηd^3^***) depends on the strength of motility. This is because the exponential component alone (-***E_A_ / E_M_***) can only report on the ratio of these two parameters. ***E_M_*** and ***E_A_*** must appear in separate fit parameters in order to infer them separately.

### Cell sorting is highly sensitive to tissue fluidity

Given the success of the minimal model in recapitulating the experimental data, we next wanted to predict how tissue fluidity is expected to influence sorting (**Fig. 2**, **Fig. 2 – Supp.** Fig. 1). In the model, one can increase tissue fluidity either by increasing the energy of cell motility (**Fig. 2a,e, Video 2**), decreasing the homotypic adhesion energy (**Fig. 2b,f**) decreasing the heterotypic adhesion energy (**Fig. 2c,g**), or by decreasing the viscosity (**Fig. 2d,h**). We therefore varied these parameters in the model to order to vary tissue fluidity (**Fig. 2e-h**) and then measured the effect on sorting after 48 hours (here and throughout, times stated refer to real time as would be measured in the laboratory, not computational time) (**Fig. 2i**-**p**). We chose 48 hours as a final timepoint because this is on the higher end of typical experimental measurement windows. Also similar to the experiments, we stopped the simulations at 48 hours regardless of whether the system reached steady state, so some measurements represent incomplete sorting. All mechanisms used to alter the tissue fluidity led to substantial changes in the degree of sorting (**Fig. 2i**-**p**), demonstrating that tissue fluidity is a critical modulator of sorting.

**Figure 2.**
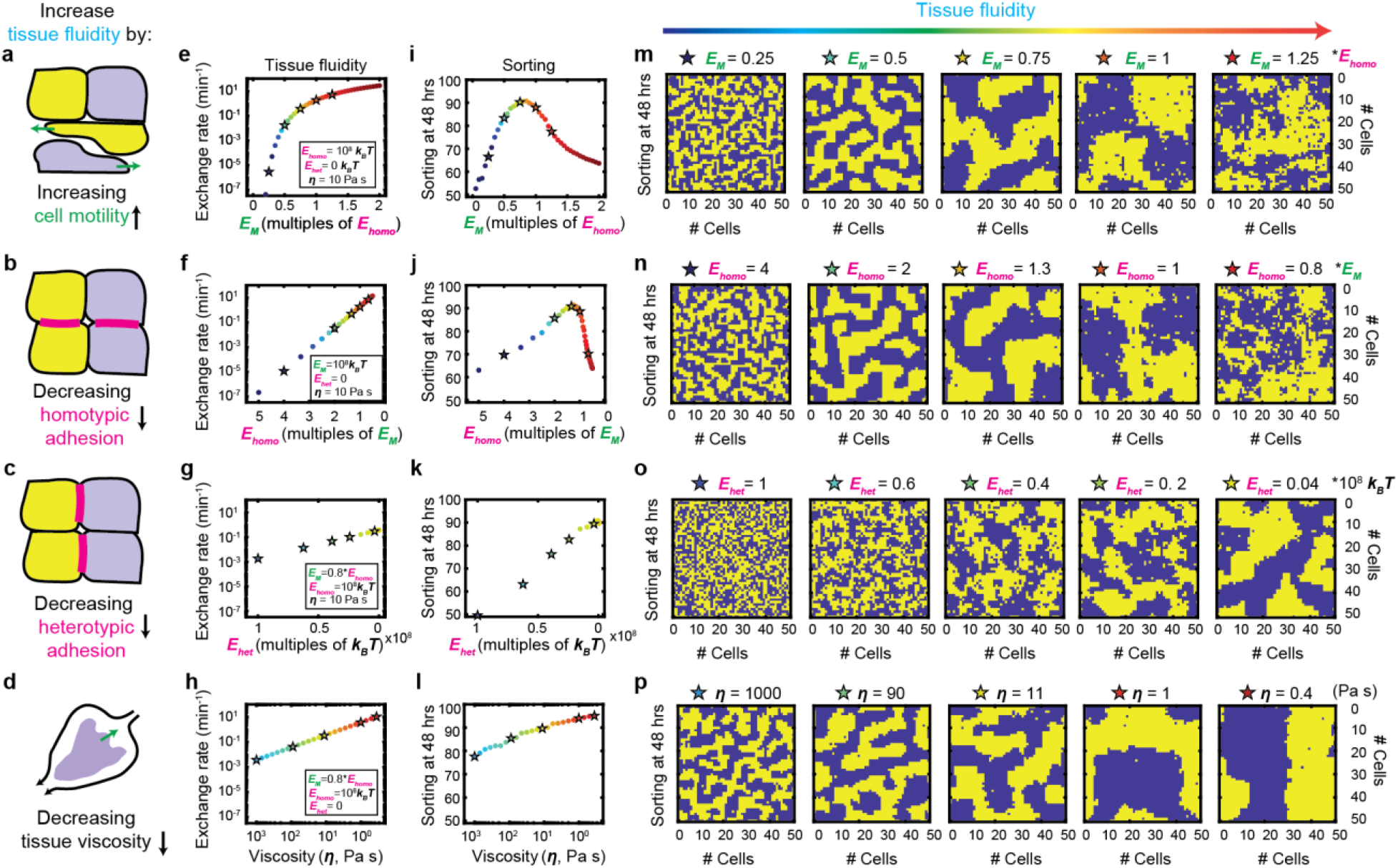
Tissue fluidity is a critical regulator of adhesion-based sorting. (**a-d**) Schematic of different mechanisms to increase tissue fluidity in the model. (**e**-**p**) Color represents tissue fluidity, where identical colormaps with identical color limits were used for (e-p). Stars in (e-h) and (i-l) represent conditions selected for display in (m-p). (**e**-**h**) Mean neighbor exchange rate at 50% sorted (two same cell type neighbors, two different cell type neighbors) as a function of the parameter varied. Choices for parameters not varied are listed in inset legend. (**i**-**l**) The degree of sorting at 48 hours (defined as the percent of each cell’s four direct neighbors (i.e., sharing faces) that are the same cell type as that cell, averaged across all cells in the tissue) is plotted as a function of the parameter varied. (**m**-**p**) The degree of sorting at 48 hours displayed as a heat map of the cell type. Note that simulations were stopped at 48 hours regardless of whether the system reached steady state.

### An intermediate tissue fluidity is optimal for sorting

Generally, increasing tissue fluidity improved sorting (**Fig. 2m [panels 1-3]**, **Fig. 2n [panels 1-3]**, **Fig. 2o-p**) by increasing the speed at which the cells sorted (**Fig. 2 – Supp.** Fig. 1, see also **Fig. 3**). Intuitively, this is because increasing tissue fluidity increases the rate of cell rearrangements within the tissue, which increases the rate at which cells of the same cell type can find each other. For the cases of heterotypic adhesion and viscosity (**Fig. 2k**,**l**,**o**,**p**), increasing tissue fluidity monotonically improved sorting—such that minimizing heterotypic adhesion and viscosity led to optimal sorting.

**Figure 3.**
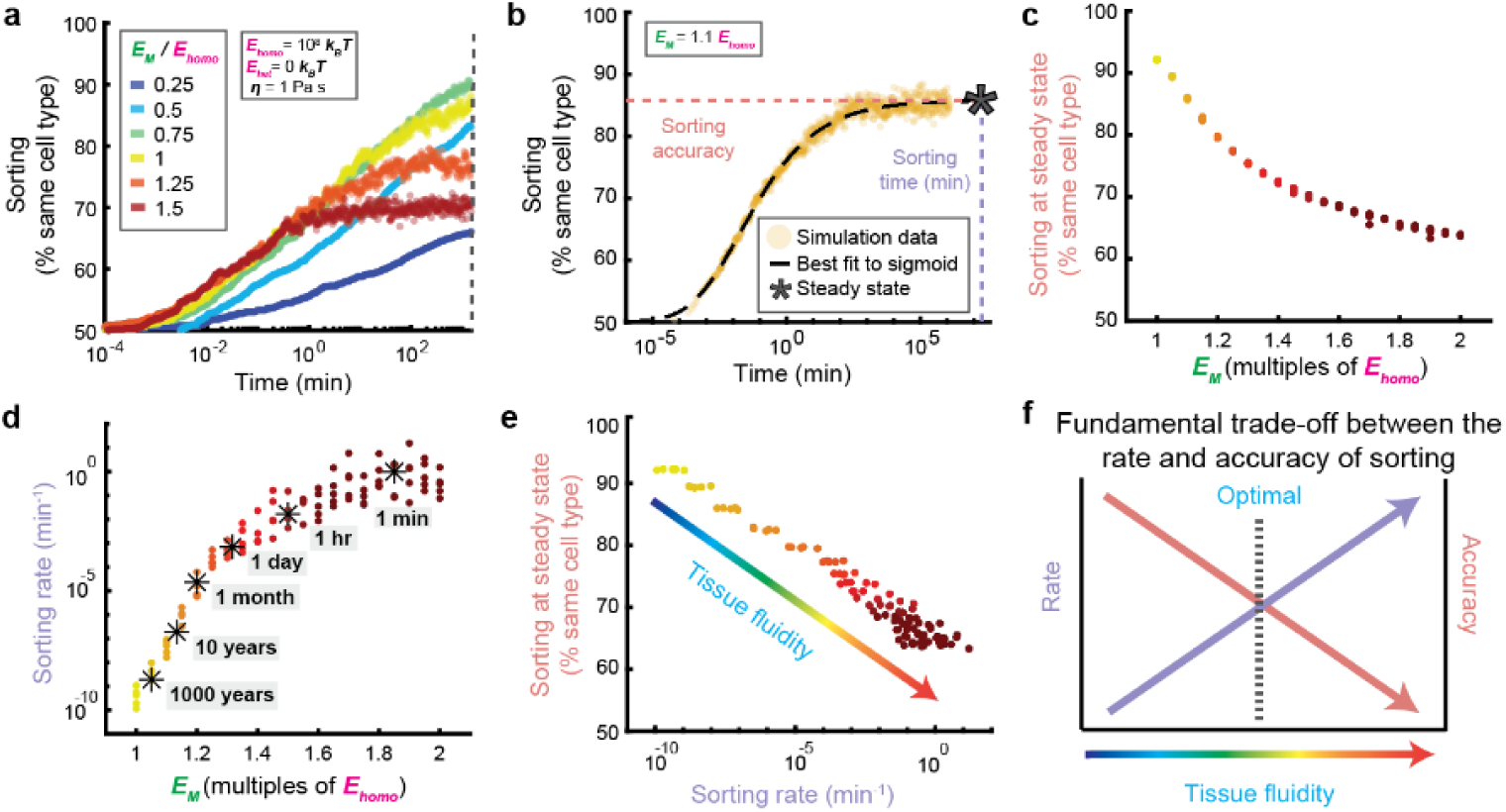
Tissue fluidity tunes a fundamental tradeoff between the sorting rate and accuracy. (**a**) Degree of sorting (i.e., average percent of a cell’s neighbors that are of the same cell type) as a function of time for simulations with variable amounts of motility energy, such that motility (and thus tissue fluidity) increase from blue to red. (**b**) Simulations were allowed to run until they reached steady state. At this point, the sorting vs time curve was fit to a sigmoid, allowing us to estimate the sorting accuracy at steady state as well as the time it took to reach steady state. The color of the curve indicates the choice of motility energy as shown in Fig. 2. (**c**) Sorting accuracy at steady state plotted as a function of the motility strength. The shape of the curve is consistent with the Boltzmann distribution shown in Fig. 3d-f. The color of each point indicates the choice of motility energy as shown in Fig. 2. (**d**) Sorting rate (defined as the inverse of the sorting time) at steady state plotted as a function of the motility strength. The color of each point indicates the choice of motility energy as shown in Fig. 2. (**e**) Sorting accuracy at steady state plotted as a function of the sorting rate for each choice of motility energy. The color of each point indicates the choice of motility energy as shown in Fig. 2. (**f**) A schematic representing the major results from (a-e). The optimal tissue fluidity arises due to a trade-off between the sorting rate, which increases with the fluidity, and the sorting accuracy, which decreases with fluidity.

However, for the case of cell motility and homotypic adhesion, an intermediate tissue fluidity was optimal for sorting. When plotting the degree of sorting after 48 hours as a function of the motility strength (**Fig. 2i**) there was a clear optimum level of motility. This optimum has also been observed in cellular Potts models of adhesion-based sorting(*40*). The optimum arose where the energy of motility was approximately equal to the cell–cell adhesion energy. Indeed, we also found that there was a corresponding optimal level of homotypic adhesion strength (**Fig. 2j**), approximately equal to the strength of cell motility. The source of this optimum became furthermore apparent when visualizing the sorting pattern and levels of cell mixing after 48 hours of sorting (**Fig. 2m,n**). Sorting initially improved with increasing levels of cell motility, but then degraded at very high levels of motility, as the pattern became dominated by noise. Most strikingly, even very small (< 2-fold) changes to the strength of motility resulted in striking changes to the overall pattern (**Fig. 2m,n**).

We note that both tissue fluidity and sorting are slightly more sensitive to changes in motility (**Fig. 2e,i**) than to changes in cell–cell adhesion (**Fig. 2f,j**) (note the difference in scale along the x-axes). This is because increasing the motility energy acts to increase tissue fluidity both by increasing the adhesion-free neighbor exchange rate (i.e., the prefactor ***k_0_ = E_M_ / 3πηd^3^***) as well as to counteract cell–cell adhesion (i.e., in the exponential ***E_A_ / E_M_***), while cell–cell adhesion appears only in the exponential. In contrast, if we were to instead assume the adhesion-free neighbor exchange rate was independent of the motility energy (a common assumption used in these types of models, which we argue is not biologically realistic), tissue fluidity and sorting would be slightly less sensitive to the motility but maintain the optimum. More specifically, varying motility would be mathematically identical to varying the inverse of adhesion (**Fig. 2f,j**).

Overall our model suggests that to achieve optimal sorting, one should minimize viscosity, minimize heterotypic adhesion, and maintain an intermediate level of cell motility and homotypic adhesion.

### The competition between cell motility and homotypic adhesion tunes a fundamental tradeoff between the sorting rate and accuracy

To understand why there might be an optimal level of cell motility and homotypic adhesion for sorting, we took a closer look at the sorting dynamics (**Fig. 3a, Video 2**). We found that at low levels of cell motility (e.g., blue curves) sorting was slow, and the rate of sorting increased as cell motility increased (from blue to red). However, at the highest levels of motility (e.g., orange and red curves), sorting increased until it stalled out at a steady state value. The degree of sorting at steady state (i.e., sorting accuracy) degraded with increasing levels of motility. Note that only the tissues with the highest two levels of motility plotted reached steady state within the 48 hour time window.

From a biological standpoint, these results can be intuitively understood based on the idea that if tissue fluidity is too low (either because the adhesion strength is too high or the motility strength is too low), cells cannot sort because they cannot move. As tissue fluidity increases, sorting proceeds more quickly. However, if tissue fluidity is too high, cells are able to escape the sorted configuration via active motility, making the pattern noisy. Overall, this analysis suggested that random cell motility and homotypic adhesion, through their effects on tissue fluidity, mediate a trade-off between the rate and accuracy of sorting.

To more quantitatively examine the apparent trade-off between the rate of sorting and the accuracy of sorting at steady state, we again varied the tissue fluidity by varying the strength of motility, but this time we ran each simulation until it reached steady state (**Fig. 3b**, see ***Methods***). By fitting the sorting dynamics to a standard function (**Fig. 3b**, **Fig. 3** - **Supp Fig. 1**, see ***Methods***), we then determined both the accuracy of sorting at steady state, as well as the time it took each simulation to reach steady state (**Fig. 3b**, see ***Methods***). The sorting accuracy at steady state decayed exponentially with the strength of motility (**Fig. 3c**) – suggesting again that lower levels of motility yield more accurate, less noisy, sorting. On the other hand, the sorting rate increased dramatically with increasing motility (**Fig. 3d**). This inverse relationship (**Fig. 3e**) represents a fundamental tradeoff between the rate and accuracy of sorting that is tuned by tissue fluidity (**Fig. 3e-f**). As a consequence, tissues with low motility are poorly sorted on biologically relevant timescales due to their slow sorting rate, and tissues with high motility are poorly sorted because their steady state sorting accuracy is low (**Fig. 2a,b,ef,i.j,m,n**, **Fig. 3a**). An intermediate motility is required to balance the trade-off between speed and accuracy.

Remarkably, we note that the time it took to reach steady state expanded by *orders of magnitude* as the motility decreased (**Fig. 3d**). Even a small (< 20%) reduction of the tissue fluidity increased the time it took to sort by 10-fold. Indeed, this dramatic dependence of the sorting time on the effective temperature (i.e., motility strength) is a known feature of this model(*39*, *43*, *46*, *47*, *61*), and has been mechanistically interpreted in the physics literature as the existence of metastable states (i.e., long lived local minima on the energy landscape)(*47*, *61*). Our results suggest that the rate of sorting is incredibly sensitive to the tissue fluidity, such that if tissue fluidity is not very tightly controlled, the tissue will not be able to sort on a timescale compatible with other biological processes such as cell fate specification and morphogenesis (**Fig. 3d**). Putting these results together, we show that tissue fluidity acts as both a powerful catalyst and unavoidable source of noise for cell sorting (**Fig. 3e-f**).

### Tissue fluidity sacrifices the sorting accuracy by favoring entropy

We next wanted to investigate why increasing tissue fluidity would decrease the sorting accuracy at steady state (**Fig. 3c**). Equilibrium statistical mechanics is often used to predict the behavior of biological systems at steady state (i.e., at equilibrium), though historically these approaches have focused on systems of interacting molecules. To predict the steady state sorting behavior of tissues, we therefore developed a tissue-scale statistical mechanics framework for adhesion-based sorting, similar to a recent analysis (*45*). In this framework, the tissue can sample different possible cell arrangements (i.e., microstates) as cells move around within the tissue (**Fig. 4a**). Depending on how well-sorted these arrangements are, the configurations will have different adhesion energies (i.e., macrostates, **Fig. 4b**). The way in which cells transition between the different configurations (i.e., energy states) can be represented using a free energy landscape (**Fig. 4c**), and the probability of being in any given state is determined by the Boltzmann distribution (**Fig. 4d-f**). For molecular systems, thermal energy is what provides the energy for molecules to switch between states. However, in the context of a tissue, actively driven cellular movements drive cell rearrangements. Therefore, we replace the typical thermal energy component of the equation (***k_B_T***) with the motility energy (***E_M_***). Our equilibrium statistical mechanics approach predicts exactly what we observed in the stochastic simulations (**Fig. 2**, **Fig. 3c**), which is that increasing the differential adhesion energy (either by increasing the homotypic cell–cell adhesion strength or by decreasing the heterotypic adhesion strength) pulls cells into the most sorted arrangement (**Fig. 4e**), while increasing the strength of motility drives cells out of the sorted state until the arrangement of cells is completely random (i.e., entropy/noise-dominated, **Fig. 4f**). Overall, we find that it is the competition between the differential adhesion energy and the strength of motility which determines the accuracy of sorting at steady state (**Fig. 4g**). Therefore, increasing the tissue fluidity either by increasing motility or decreasing homotypic cell– cell adhesion will always sacrifice sorting accuracy at steady state. We note that while this statistical mechanics approach allows us to analyze steady states analytically, it ignores the kinetics, which as the stochastic simulations show, vary substantially across parameter space.

**Figure 4.**
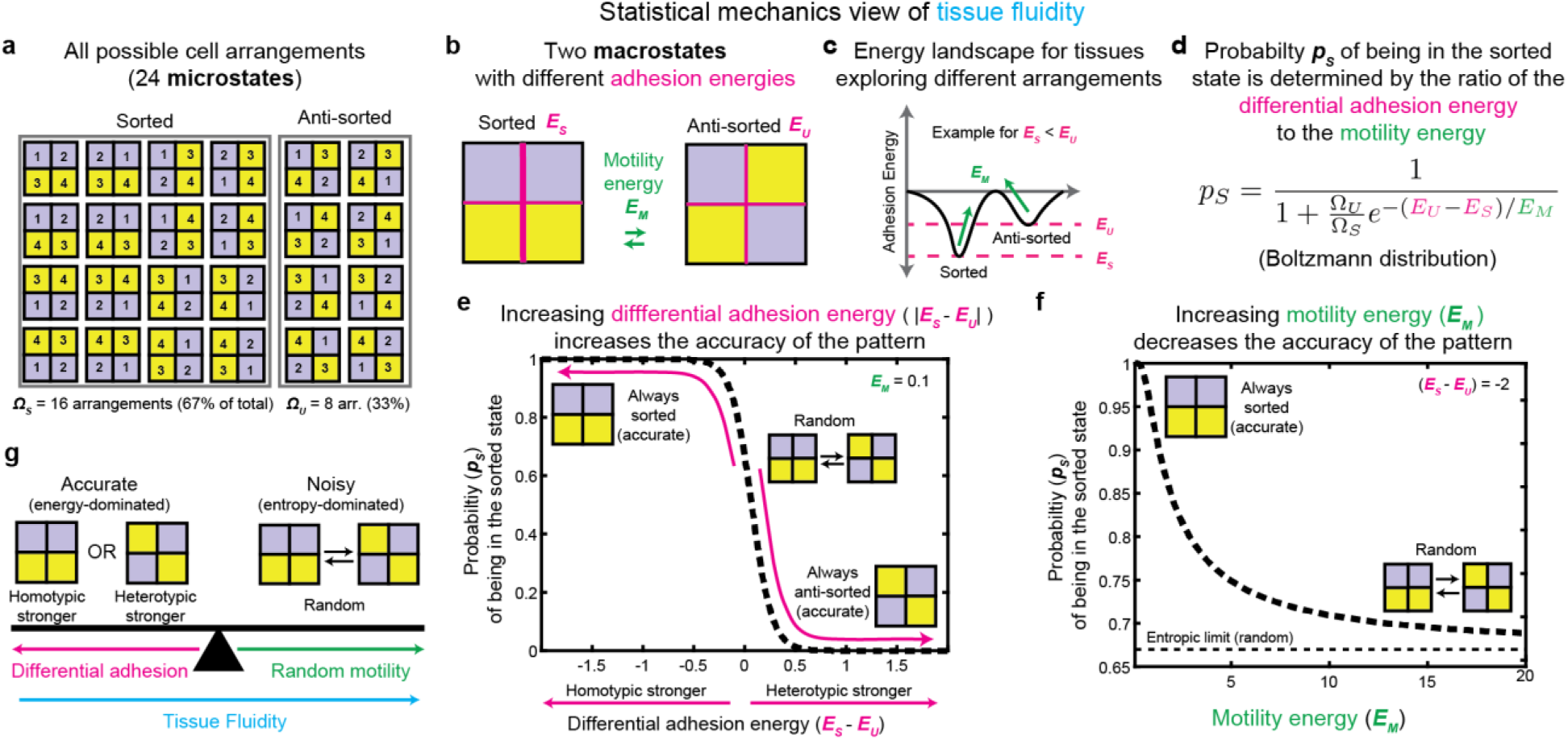
Tissue fluidity sacrifices the sorting accuracy by favoring entropy. (**a**) For an example tissue containing 4 cells of two different cell types, all possible arrangements of the 4 cells are shown. In an equilibrium statistical mechanics framework, each of these arrangements represents an individual microstate. 16 of these microstates are in the sorted configuration (where cells of the same cell type are adjacent), and 8 of these microstates are anti-sorted. (**b**) Because the sorted and anti-sorted configurations have different numbers of homotypic and heterotypic cell–cell interfaces, they each have different adhesion energies. In an equilibrium statistical mechanics framework, the sorted and anti-sorted configurations represent individual macrostates. (**c**) The way in which cells transition between the sorted and anti-sorted energy states can be represented as a free energy landscape. The sorted and anti-sorted states represent energy wells, where the adhesion energy of each state represents the energy barrier that must be overcome to switch between states. The motility strength provides the energy for cells to overcome this energy barrier. For the case shown, the homotypic energy is stronger than the heterotypic adhesion energy, so the sorted state lies in a deeper well. (**d**) For a system at thermodynamic equilibrium, the Boltzmann distribution describes the probability that a system will be in a given state, based on the energy of each state, the number of microstates in each state, and the energy available to escape any given energy state. The probability of being in the sorted state increases with the differential adhesion energy between the two states, and decreases with increasing motility. (**e**) Plotting the equation shown in (d): The probability of being in the sorted state as a function of the differential adhesion energy. The stronger the differential adhesion energy, the more likely the tissue will be in the sorted state (if homotypic adhesion is stronger) or in the in the anti-sorted state (if heterotypic adhesion is stronger). The smaller the differential adhesion, the more likely the system will rapidly switch between either state at random. (**f**) Plotting the equation shown in (d): The probability of being in the sorted state as a function of the motility energy. The stronger the motility energy, the more likely the system will rapidly switch between either state at random. (**g**) Schematic of the results shown in (e-f). A competition between differential adhesion and random motility determines whether the tissue will be stably in either the sorted or anti-sorted state (energy dominated) or whether the tissue will rapidly oscillate between states at random (entropy dominated). Although tissue fluidity is set by the competition between total adhesion (not differential adhesion) and motility strength, the total adhesion typically also increases with the differential adhesion. Therefore, tissue fluidity will typically increase from left to right.

Taken together, we find that the glassy dynamics of biological tissues require high effective temperatures (i.e., high motility), approaching the critical temperature(*61*), in order for phase separation to occur on a biologically-relevant timescale. However, as the system approaches the critical temperature, it quickly encounters an entropy dominated regime where random mixing dominates over sorting(*65*, *66*). This puts tissues in stark contrast to many molecular phase separating systems such as lipids and biomolecular condensates, which do not typically exhibit these glassy dynamics because they are much smaller, experience lower viscous drag, and therefore diffuse orders of magnitude faster. Molecular systems thus sort quickly at temperatures well below the critical temperature, and consequently do not exhibit this trade-off(*51*, *65*). Overall, we find that the unique properties of biological tissues mandate that cell sorting sits between metastable glassy regime (a “rock”) and an entropy-dominated regime (a “hot place”), with an intermediate motility balancing this trade-off.

### Sorting requires a tight coupling between motility and adhesion

The results thus far suggest that tissue fluidity must be tightly controlled to maintain the ability of cells to sort. The optimal fluidity lies where the motility energy is approximately equal to the adhesion energy (**Fig. 2i-j**, **Fig. 5a**). However, the coupling between the motility and adhesion required to carry out sorting is so tight that even a two-fold change in either parameter can dramatically reduce sorting (**Fig. 5a**), whereas back-of-the envelope calculations for the adhesion and motility energies suggested these values can both vary widely under different conditions (**Fig. 5b**). This raises the question of how cell sorting can be robust to fluctuations in adhesion protein expression given this constraint.

**Figure 5.**
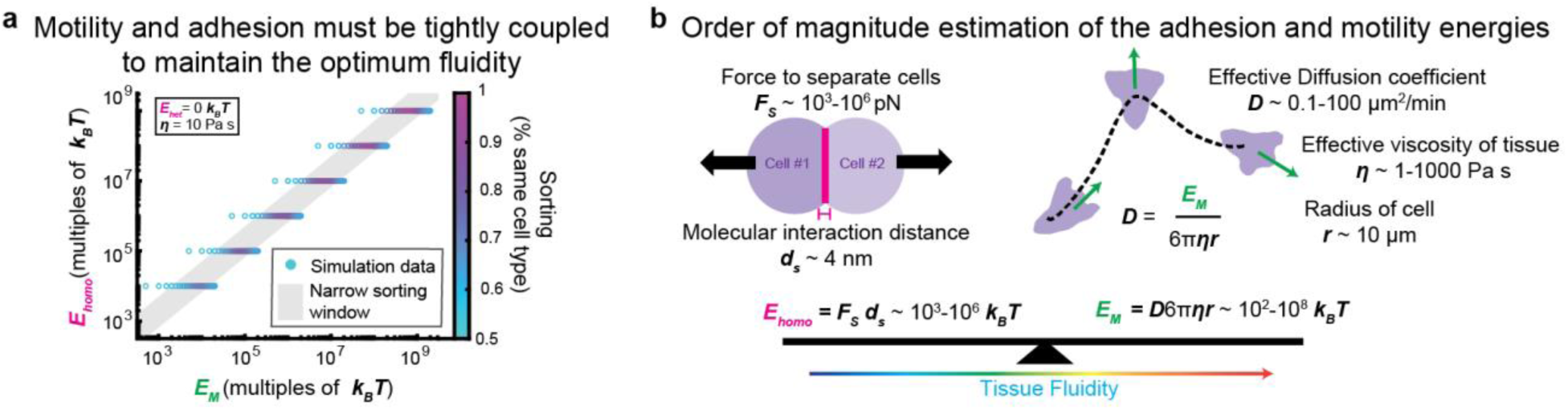
Sorting requires a tight coupling between motility an adhesion. (**a**) The degree of sorting after 48 hours of sorting time is plotted as a function of the choices for motility energy and adhesion energy in the simulation. Sorting can only occur in a very narrow window where the adhesion and motility energies are nearly equal. (**b**) Schematic of the order of magnitude estimation for the values of the adhesion (left) and motility (right) energies. *Left:* The adhesion energy is defined as the energy required to separate two adherent cells. This energy can be estimated as the force required to separate two adherent cells (e.g., as measured by pipette pulling experiments) multiplied by the distance the cells must be separated by before the cadherin molecules are no longer in contact. *Right:* The motility energy is calculated as the effective diffusion coefficient of motility (e.g., as measured by cell tracking) multiplied by the viscous (Stokes’) drag coefficient for a cell as it migrates through the tissue.

To determine how sensitive sorting is to tissue fluidity, we returned to our experimental cell sorting assay (**Fig. 1g**). To vary the fluidity experimentally, we opted to change the cell–cell adhesion strength. In particular, we generated four L929 cell populations with high or low expression of either *Cdh2* (N-cadherin), or *Cdh3* (P-cadherin), and mixed them with cells expressing either high or low expression of *Cdh1* (E-cadherin), respectively. As before, the *Cdh2*-and *Cdh3*-expressing cells also co-expressed *EGFP* from the same expression cassette separated by a P2A self-cleaving peptide linker, and the *Cdh1*-expressing cells co-expressed *mRFP1* in an equivalent manner. The high- and low-expressing populations exhibited a roughly 2-fold difference in average fluorescence intensity (**Fig. 6 – Supp.** Fig. 1) when imaged under the same conditions. We then performed time-lapse imaging of cell sorting under these conditions, and quantified the sorting dynamics.

**Figure 6.**
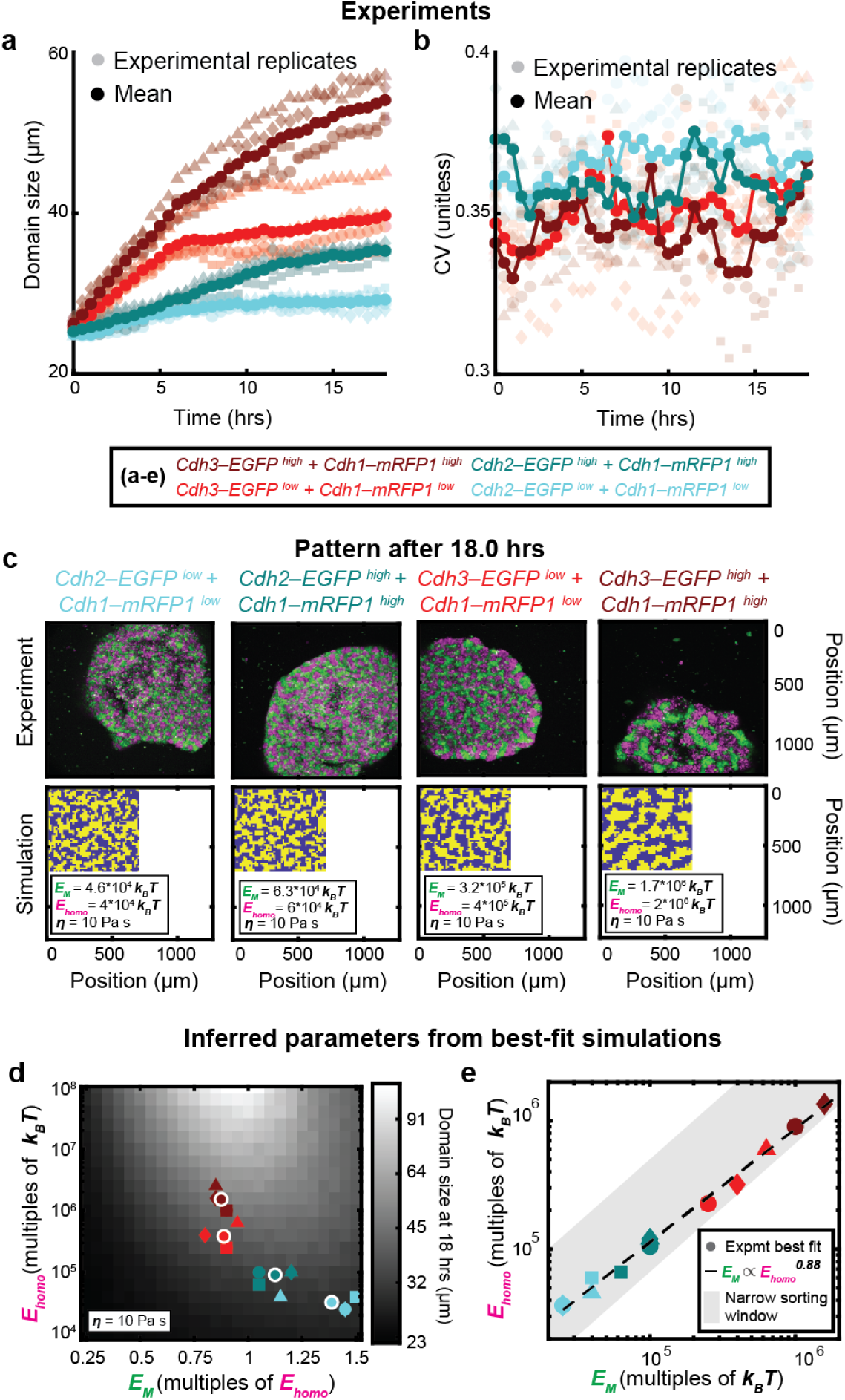
Cells exhibit strong apparent coupling of motility and adhesion. (**a-b**) The average (a) and coefficient of variation (b) of the domain size are plotted as a function of time for each of the 4 different experimental conditions. Light markers: Replicates. Dark Markers: Average across 4 replicates. Sorting can only occur in a very narrow window where the adhesion and motility energies are nearly equal. (**c**) Representative images of each experimental cell-sorting assay (*top*) and its associated representative best-fit simulation (*bottom*) after 18 hours. *Top:* L929 cells co-expressing either high or low levels of *Cdh2* (N-cadherin) or *Cdh3* (P-cadherin) and *EGFP* (green) were mixed in equal proportions with cells co-expressing either high or low levels of *Cdh1* (E-cadherin) and *mRFP1* (magenta) and imaged by confocal time-lapse microscopy. Images represent maximum intensity projections. *Bottom:* Best-fit simulations with displayed as a heat map of cell type. Best fit parameters are listed in the inset legend. (**d**) The best fit simulation parameters for each experimental dataset are plotted on top of a heatmap of the average domain size for all simulation parameters tested. Markers with a white outline represent the mean across the 4 replicates for each condition. (**e**) The best fit adhesion energy is plotted as a function of the best fit motility energy for each experimental dataset. The narrow sorting window is plotted exactly as in Fig. 5a.

Each of the four experimental conditions (*Cdh2–EGFP*^high^ cells with *Cdh1–mRFP1*^high^ cells, *Cdh2– EGFP*^low^ cells with *Cdh1*–*mRFP1*^low^ cells, *Cdh3–EGFP^high^* cells with *Cdh1–mRFP1^high^* cells, and *Cdh3*–*EGFP^low^* cells with *Cdh1*–*mRFP1^low^*cells) exhibited some amount of sorting. Within each combination of cadherin identities, the high-expressing combination sorted better than the low-expressing combination, as expected. In addition, conditions in which *Cdh3-* and *Cdh1-* expressing cells were mixed (**Fig. 6a**, red) consistently yielded a greater degree of sorting, as measured by average domain size, than those in which *Cdh2*- and *Cdh1*-expressing cells were mixed (**Fig. 6a**, cyan). Across all datasets, there was a general trend wherein conditions with higher levels of sorting tended to exhibit slightly less noisy patterns, as measured by the coefficient of variation (CV) of the domain size (**Fig. 6b**).

For each of these conditions, we again performed automated comparisons between the sorting dynamics in the experiments and the simulations (**Fig. 6c-e**, **Fig. 6 – Supp.** Fig. 2, **Fig. 6 – Supp.** Fig. 3), to determine the best fit simulations parameters for each experimental condition (**Fig. 6c-d**). Again, this fitting approach allows us to estimate the values of the motility energy (***E_M_***) and homotypic adhesion energy (***E_A_***), assuming a given choice of viscosity (***η***) – and requires that the model’s adhesion-free neighbor exchange rate (***k_0_ = E_M_ / 3πηd^3^***) depends on the strength of motility, ***E_M_***. Just as for the original assay (**Fig. 1g-i**), the best-fit simulations for each experimental condition were able to recapitulate the experimentally-observed patterns, as well as the growth dynamics of the mean and standard deviation in the domain size over time (**Fig. 6 – Supp.** Fig. 2). Heatmaps of the cost function used for fitting show that the best fit simulation parameters occupied reasonably well-defined minima in parameter space (**Fig. 6 – Supp.** Fig. 3). The clustering of the of the best fit parameters for the experimental replicates also suggests that these values are fairly well-constrained (**Fig. 6d**).

The best fit adhesion energies roughly followed the adhesion protein expression levels as measured by the fluorescence intensity, such that the difference in intensity was approximately 2-fold between low and high expressing conditions, and the difference in inferred adhesion energy was 1.5-fold for the conditions mixing *Cdh2–EGFP* cells with *Cdh1–mRFP1* cells, and 5-fold for the conditions mixing *Cdh3–EGFP* cells with *Cdh1–mRFP1* cells. This agreement is reasonable given the known nonlinearity between adhesion protein expression and adhesion strength(*67*). However, we stress that this minimal tissue model is not able to account for effects such as differences in tissue compaction between experimental conditions, changes in adhesion protein expression over time, or more complex cell deformations and movements, which could lead to errors in estimates of the adhesion and motility energies. Further, the model assumes the homotypic adhesion energies of the two cell types being mixed were identical, so when two cell types with different homotypic adhesion energies are mixed in experiments, the best fit model returns a single value representing an approximate average adhesion energy in the tissue. Finally, the choice of a square grid assumes each cell has four neighbors, which will lead to a small, systematic offset in the fit estimates for tissues with a different number of neighbors (see ***Methods,* Fig. 6 – Supp.** Fig. 4).

To understand how cell motility and cell–cell adhesion might be coupled in living cells, we plotted the best fit adhesion energy as a function of the best fit motility energy across all experimental conditions (**Fig. 6d-e**). Interestingly, we find that the strength of cell motility appears to increase monotonically with the cell–cell adhesion, precisely following a scaling law of 0.88 (**Fig. 6e**). While any experimental result that yields sorting must fall within the gray sorting window (**Fig. 6e**), the precise power law scaling suggests a tight coupling between these two physiological parameters. These results provide one potential explanation for how sorting can occur despite the narrow constraints on tissue fluidity – suggesting that cells may not tune motility and adhesion strength independently, but rather may naturally scale their motility with their adhesion strength to maintain a permissible fluidity.

## Discussion

### Co-evolution of tissue fluidity with patterning mechanisms

Overall, our results suggest that (1) tissue fluidity must be tightly regulated to carry out sorting that is both fast and accurate, and (2) a moderate amount of cell rearrangement is absolutely required for sorting to occur. It is interesting to contrast adhesion-based sorting to the classic French Flag morphogen gradient mechanism of patterning, which in its original formulation requires a completely solid tissue (stationary cells) to achieve optimal patterning.

Intriguingly, levels of tissue fluidity vary drastically across species, tissues, and developmental time. In some organisms, such as worms, flies, frogs, and ascidians, development is fairly deterministic, such that there is little to no random mixing of cells during embryogenesis(*69–72*). On the other hand, random motility, active cell mixing, and scattering of clonal cell lineages have frequently been observed in zebrafish(*73–75*), chick(*36*, *76*, *77*), mouse(*24*, *78–82*), and human(*83*) embryos.

It is hard to imagine how adhesion-based sorting could work in an organism with no cell mixing, or how a French-flag type morphogen gradient mechanism could operate in a tissue with extensive cell rearrangement. Together, these observations suggest that patterning mechanisms must have had to co-evolve with tissue fluidity to achieve optimal patterning in different developmental contexts.

### Molecular and cellular mechanisms regulating tissue fluidity

Another major conclusion of our work is that living cells may intrinsically co-regulate cell motility and adhesion in order to maintain an optimal level of tissue fluidity. The observed power law scaling between these two properties in the experimental assay suggests this behavior may be driven by a fundamental biophysical relationship (*84–86*). What might be the molecular and cellular mechanisms that allow the strength of cell motility and cell–cell adhesion to be so tightly coupled? We speculate that the answer might be biophysical in nature. At the molecular level, the branched actin networks that drive leading edge protrusion are remarkably responsive to obstacles or other resistive forces (*87*, *88*). These networks naturally increase their actin polymerization rate when they encounter a barrier, allowing them to increase their protrusive strength and push back proportionally (*87*, *88*). It is possible that in cases of high cell–cell adhesion, the cytoskeleton naturally polymerizes more actin at the cell’s leading edge to compensate for the increased resistance to movement provided by cell–cell adhesion.

Another possibility is that cells migrate more efficiently in the context of higher cell–cell adhesion. While perhaps counter-intuitive, cells in a confluent tissue are essentially crawling on each other, and so therefore may have more traction (i.e., push off of each other with less slipping) in the context of high cell–cell adhesion—in the same way it would be easier to shimmy up a sticky chimney than a slippery one. This plays out at the molecular level in that higher cell–cell adhesion could more solidly couple the F-actin cytoskeleton to the cell’s extracellular environment, such that there is less slipping (i.e., retrograde flow) of actin filaments and therefore more efficient protrusion.

On the other hand, the feedback could occur at the level of signaling or gene expression. β-catenin, a protein which physically links cadherins to the actin cytoskeleton, also acts as a transcription co-regulator upstream of motility-related genes. It is possible that increases in adhesion molecule expression lead to an increase in motility via its modulation of β-catenin transcription factor activity. Additionally, cadherins are capable of remodeling the actin cytoskeleton and vice versa(*89*). This coupling of actin organization and cadherin clustering and turnover dynamics could also be responsible for the robust scaling. The experimental generality of the coupling of motility and adhesion energies as well its molecular underpinnings will require further investigations.

### Biological complexity: Additional mechanisms driving tissue fluidity

Many different kinds of models have been developed to investigate tissue fluidity and/or cell sorting(*41*), including lattice models(*39*, *43*, *45*), finite-element and finite-volume models(*90–92*), cellular Potts models(*22*, *23*, *40*, *42*, *93*, *94*), vertex models(*37*, *38*, *44*, *54*, *55*, *60*, *95–98*), active foam models(*35*, *53*), and self-propelled particle models(*46*). The main advantage of our lattice modeling approach is its conceptual simplicity and computational speed, allowing us to rapidly and efficiently fit the model to experimental data across large parameter sets. Moving forward, it will be interesting to explore these same concepts in models of tissue fluidity with sub-cellular resolution (e.g., cellular Potts, vertex, and active-foam models), which are able to directly incorporate dynamics in cell shape and cell–cell interactions. While our model considers the roles of cell motility, cell–cell adhesion, and viscosity in tuning tissue fluidity, these other models excitingly provide additional means by which to tune tissue fluidity, for example by altering the stiffness of the cell cortex(*54*), or fluctuations in contractility along cell–cell interfaces(*47*, *53*). Indeed, vertex models have shown that, similar to motility as used in this model, active junctional fluctuations at cell–cell interfaces can also speed the sorting rate(*47*). Additionally, self-propelled particle models have shown that coupling of cell motility between neighboring cells significantly reduces the amount of time it takes to sort(*46*), potentially allowing tissues to reduce the steep the trade-off between the rate and accuracy of sorting reported in this work. It will be fascinating to see how these other fundamental biophysical properties can be tuned to optimize tissue patterning.

Even within the current modeling approach, there are many exciting future directions regarding how tissue fluidity might interact with more complex aspects of biological cell sorting. While the current work focused on the reticulated patterns observed in the experimental system (i.e., those that arise from equal homotypic adhesion energies and limited heterotypic adhesion energy), other types of patterns produced in different regimes, such as inside-outside patterns and checkerboard patterns, may be more or less sensitive to tissue fluidity. A checkerboard pattern, for example, requires less neighbor exchange than patterns with larger domains, and so may require lower tissue fluidity to achieve accurate sorting. The model could also be used to explore trade-offs with tissue fluidity in the context of unequal or rare cell type population frequencies – where a rare cell type may require higher tissue fluidity to search over larger distances and find its partners. One could also investigate how the optimal tissue fluidity varies under conditions including a larger diversity of cell types, cell-type-specific differences in motility, time-varying parameters, and feedback between parameters – indeed, many these complexities have been explored in related modeling approaches(*19*, *21*, *23*, *27*, *40–42*, *47*, *63*, *90*). Additionally, the incorporation of moving boundary conditions could enable insights into more dynamic tissues that exhibit burst-like spreading and asymmetric domain expansion, where clear feedbacks exist between cell–cell adhesion, motility, and tissue boundaries(*64*). This approach could also be extended to different lattice geometries (*68*, *99*).

In at least two other models, motility was also treated as an effective temperature(*40*, *45*). Agreeing with our findings, cellular Potts modeling has also demonstrated that there is an optimal level of motility for sorting(*40*). However, the authors of this previous work attributed the optimum to sorting being fastest at an intermediate motility: “Cell motilities above or below the optimum sort more slowly”(*40*). In contrast, our work shows that the sorting rate monotonically increases with the level of cell motility, and the poor sorting at high levels of motility arises from poor sorting accuracy at steady state (rather than a slow-down in the sorting rate). Indeed, our finding agrees with recent theoretical and experimental work suggesting that cell motility drives structural heterogeneity in tissue patterning(*45*).

### Model adaptability: Exploring the role of tissue fluidity across diverse patterning mechanisms

Our work shows that tissue fluidity is an important factor in controlling adhesion-based sorting, and is likely to be important for other patterning mechanisms as well. A major advantage of our minimal modeling approach is that it can easily be integrated with different types of biochemical signaling (e.g., morphogen gradient signaling, Notch-Delta signaling), to explore how they interplay with tissue fluidity to control cell patterning. Given the sensitive way in which adhesion-based sorting responds to tissue fluidity, it seems likely that the optimal tissue fluidity will vary for different patterning mechanisms. In addition, it is often the case in embryonic development that multiple different mechanisms act in synchrony to pattern tissues(*16*, *23–25*, *36*, *100*, *101*). Our modeling approach provides a path forward towards understanding how tissue fluidity controls patterning across these diverse developmental contexts.

## Methods

### Model description

All modeling and model analysis were performed in MATLAB.

#### Cell and tissue geometry

Tissues were modeled as a square grid of cells. The grid represents a confluent tissue (i.e., no holes), wherein cells can only move within the tissue by swapping places with one another (i.e., neighbor exchange). By default, the model included a 50x50-cell grid with periodic boundary conditions, where neighbors were defined as any two cells sharing faces (but not corners, i.e., 4-pixel connectivity). Because the tissue is assumed to be confluent, the distance between cell centers, ***d***, is assumed to be twice the cell radius ***r***. such that ***d***=2***r***. By default, a cell radius ***r***=7µm was used, which was chosen to be roughly comparable to the cell radius in the experiments (data not shown).

We chose a square lattice because the experimental system did not display an obvious hexagonal arrangement of cells, and therefore we chose the simplest possible grid arrangement for the model. Previous work on the coarse-grained Kawasaki model has shown that changing the number of neighbors (i.e., coordination number of the lattice) results in a rescaling of the sorting timescale, without affecting the qualitative dynamics(*68*). Indeed, using simulations, we show that the sorting timescale differs by approximately 4-fold between a square and hexagonal lattice (**Fig. 6 – Supp.** Fig. 4). In terms of fitting the model to the experimental data, different lattice geometries will yield small multiplicative differences in the inferred magnitude of the motility and adhesion energies, but should not affect the ratio of these two parameters.

#### Motility energy

The strength of motility for any individual cell was modeled as an effective thermal energy, ***E_M_***. By default, all cells were assumed to have the same motility strength, and the motility strength was assumed to be constant in time.

To get an order of magnitude estimate of the motility energy, we assumed the random motility of cells within a tissue can be modeled as Stokes-Einstein diffusion at low Reynolds number (Fig. 5b), such that the diffusion coefficient ***D=E_M_***/***γ*** is equal to the motility energy ***E_M_*** divided by the viscous drag coefficient ***γ*** (see below). To estimate the diffusion coefficient for random migration, we looked at observed rates of cell migration in tissues. The fastest migrating cells in the body, such as neutrophils, can migrate their body length (∼10 µm) in one minute (*102*), giving a maximum diffusion coefficient of 100 um^2^/min. The lower bound of the diffusion coefficient is of course 0, however a reasonable cut-off for the slowest movement that would be considered migration for the purposes of sorting would be to move the same distance (10 µm) in one day, or ∼0.1 um^2^/min. Plugging these minimum and maximum estimations for the cell diffusivity as well as the viscosity (see below) into the equation ***E_M_=Dγ***, we arrive at an estimate of the motility energy ***E_M_*** between 10^2^-10^8^ ***k_B_T***.

#### Adhesion energy

At each time step, each cell–cell interface of all possible neighbor pairs was assigned an adhesion energy, ***E_I_***, based on whether the two cells were the same cell type (homotypic) or different cell types (heterotypic). The homotypic and heterotypic adhesion energies, ***E_homo_*** and ***E_het_***, were assumed to be constant in time. By default, heterotypic adhesion energy was 0. The total adhesion energy holding each cell in place, ***E_T_*** = ***N_homo_E_homo_ + N_het_E_het_***, was defined as the sum of the adhesion energies across all of the cell’s interfaces with its neighbors. To determine the neighbor exchange rate for a neighbor pair, the sum of the total adhesion energy experienced by both cells in the pair, ***E_A_= E_T_^1^+E_T_^2^***, was used.

To get an order of magnitude estimate of the adhesion energy, we took two separate approaches. First, we reasoned that the adhesion energy at a single interface between two cells ***E_homo/het_*** = ***F_s_d_s_*** should be equal to the force required to separate the cells, ***F_s_***, multiplied by the distance at which the force must be applied until the cells come apart, ***d_s_***. Therefore, we took experimental dual pipette measurements of the force required to separate two cells in a cell doublet pair, which is typically ∼ 10^0^-10^3^ nN(*16*, *103*), and multiplied these estimates by the typical overlap distance between cadherin molecules interacting in *trans*, which is a few nanometers (*104*), giving an estimate of the adhesion energy ***E_homo/het_*** between 10^3^-10^6^ ***k_B_T***.

As an alternative estimate of the adhesion energy, we reasoned that the adhesion energy at a given cell–cell interface ***E_homo/het_*** = ***E_B_N_B_*** should be the energy of any given cadherin bond, ***E_B_***, multiplied by the number of cadherin bonds at a typical cell–cell interface, ***N_B_***. Based on measurements from analytical ultracentrifugation and surface plasmon resonance (*105*) as well as force pulling experiments (*106*), the energy of a typical single cadherin binding interaction is on the order of ∼10 ***k_B_T***. Based on super-resolution imaging of E-cadherin in cells, the number of cadherin molecules at a single cell–cell interface ***N_B_*** is on the order of 10^3^-10^6^. Combining these estimates, we arrive at an estimate of the adhesion energy ***E_homo/het_*** between 10^4^-10^7^ ***k_B_T***.

#### Viscous drag

The viscous drag on the cell was assumed to be Stokes’ drag, such that the viscous drag ***γ = 6πηR***, is determined by the viscosity, ***η***, and the cell radius, ***R***. The viscosity represents the effective viscosity experienced by the cell during cell rearrangement due to effects other than adhesion (e.g., viscoelastic cell deformation, friction between neighbors, extracellular matrix, extracellular fluid viscosity). Increasing viscosity reduces tissue fluidity. The viscosity of water sets a lower bound for the viscosity at 10^-3^ Pa s, or 10^-6^ pN ms nm^-2^. In the early zebrafish embryo, the tissue-level viscosity of the blastoderm was measured by micropipette aspiration to be on the order of 10^0^-10^3^ Pa s depending on the stage of development (*25*), and higher tissue viscosities have been reported, so a reasonable viscosity ***η*** could vary anywhere within the range of 10^-3^ to 10^3^ Pa s. However, most reported tissue viscosities are measured in the context of tissues with non-zero cell–cell adhesion, and so are likely over-estimates of the adhesion-free viscosity used in this model. In reality, the adhesion-free viscosity is likely somewhere in between the viscosity of the extracellular fluid (e.g., water ∼10^-3^ Pa s), and the viscosity of the cell cytoplasm or cortex (∼ 100 Pa s).

#### Neighbor exchange rate

At each time step, the rate of neighbor exchange, ***k***, was calculated for all possible neighbor pairs, based on the total adhesion of each cell with its neighbors, ***E_A_***, the cell motility, ***E_M_***, the viscous drag on the cell, ***γ***, and the distance between cell centers, ***d***, – following the Arrhenius relationship ***k*** = ***k_0_ exp(***–***E_A_/E_M_)***, with a prefactor ***k_0_*** = ***E_M_ / γd^2^*** (Fig. 1c).

#### Neighbor exchange probability

At each time step, the probability of neighbor exchange, ***p***, was calculated for all possible neighbor pairs, assuming that neighbor exchange is governed by Poisson statistics. The probability of neighbor exchange ***p = k_*_dt*** was thus defined as the rate of neighbor exchange rate multiplied by the duration of the time step.

#### Neighbor exchange implementation

At each time step, a random number generator assigned each cell pair a random number from 0 to 1. If the randomly assigned number was less than the probability of a neighbor exchange, ***p***, the neighbors were swapped. If not, the cells remained in their initial positions.

#### Time steps

At each time step, the duration of the time step was set sufficiently small that a single cell should not swap with more than one neighbor within the time step. This time step duration was set automatically based on the highest neighbor exchange rate in the tissue at that time step. An additional check was performed at each iteration to ensure that a single cell never swaps with more than one neighbor in any given time step. In the case that a single cell was chosen to swap with more than one neighbor, then the time interval was dropped by a factor of two and all calculations for that time step were repeated.

#### Simulation duration

For simulations run for a fixed amount of time, the simulation continued until the last set time point was recorded. For simulations run to steady state, see below.

### Domain size scaling

Classic Kawasaki Ising behavior is characterized by a power law scaling of the domain size with t^1/3^. We therefore analyzed the scaling of the domain size with time in our model (DT local, **Fig. 1 – Supp.** Fig. 2a-e), and found that sorting exhibited sub- t^1/3^ growth dynamics. To understand and confirm this nonclassical behavior, we generated two other versions of the Ising Kawasaki model using Monte Carlo simulations implemented via a Metropolis algorithm. In one version, only directly adjacent neighbors could swap within a given Monte Carlo step (MC local, **Fig. 1 – Supp.** Fig. 2e), representing intermediate-time behavior wherein cells can exchange places with their neighbors sufficiently quickly to find the lowest energy configuration within a time step. In the other version, any two cells on the grid could swap (MC global, **Fig. 1 – Supp.** Fig. 2e), representing long-time behavior wherein cells can diffuse across the grid sufficiently quickly to sample large displacements within a time step. Only the Monte Carlo implementation where any cells on the grid can swap exhibited the t^1/3^ power law scaling (MC global, **Fig. 1 – Supp.** Fig. 2e). The divergence from the t^1/3^ power law scaling in the other two models, which are expected to more accurately resolve behavior at shorter timescales, demonstrates that sorting of cells within tissues on biologically-relevant timescales occurs well within the short-time glassy regime(*61*), long before the system reaches the t^1/3^ scaling regime.

Each of the models exhibited different dependences on the motility energy. At high temperatures, all three model implementations converge (**Fig. 1 – Supp.** Fig. 2f). This convergence is expected given that the system is well-mixed within this regime, thus satisfying the well-mixed assumption of the Monte Carlo model. It also provides confidence in the accuracy of the discrete time implementation. Therefore, our fixed time step implementation converges with the Monte Carlo models in the high temperature regime where the models are expected to be equivalent, and provides a more accurate representation at low temperatures where the assumptions of the Monte Carlo modeling approach are no longer valid.

### Fitting the sorting rate and accuracy in simulations run to steady state

For each simulation run to steady state (**Fig. 3**, **Fig. 3** - **Supp Fig. 1**), the degree of sorting (average fraction of each cell’s neighbors that are the same cell type, which ranges from 0.5 to 1 for cases where homotypic interactions are preferred) was recorded at regular intervals. The recording intervals were uniformly spaced on a log scale. Every 300 timepoints, a check was automatically performed to determine whether the simulation had reached steady state.

To determine whether the simulation had reached steady state, the sorting vs time curve was fit to an asymmetric Hill function (**Fig. 3b**): ***S***(***t***) = 0.5+(**a**-0.5)/(1+((***b***/***t***)^***c***))^***d***, where ***S***(***t***) is the degree of sorting as a function of the time, ***t***. The four best fit parameters are ***a***, the degree of sorting at steady state (i.e., sorting accuracy), ***b***, the characteristic time scale, ***c*** is the Hill coefficient (steepness), and ***d***, the asymmetry of the curve. From this best fit curve, the sorting time was defined as the time, ***t_SS_***, when the predicted degree of sorting was within 0.001 of the sorting accuracy at steady state ***a*** (i.e., when |***S***(***t***)-***a***|<0.001).

If two conditions were met, then the simulation was considered to have reached steady state and was terminated. If the conditions were not met, the target total simulation time was increased by a factor of 2, and the check was performed after another 300 timepoints. The conditions were that (1) the simulation time had to be larger than the predicted time to reach steady state ***t_SS_***, and (2) the time-averaged, cell-averaged degree of sorting over the last 21 timepoints had to be at least one standard deviation from the predicted sorting accuracy at steady state.

Once the simulation had reached steady state, the sorting vs time curve was once again fit to the asymmetric Hill function, where the sorting accuracy at steady state was defined as the best fit parameter ***a*** (**Fig. 3b,c,e**), and the time to reach steady state was defined as ***t_SS_*** (**Fig. 3b,d,e**).

### Pre-processing of the simulation data for domain size calculations

*Cell type classification –* The simulations generated cell type classifications by default (a 2D grid of cells where each pixel represents one cell, and where cells of cell type #1 were labeled with a 1 and cells of cell type #2 were labeled with a 2).

#### Up-sampling to experimental resolution

In the simulations, each pixel represents one cell. In the experiments, a single cell contained many pixels. In order to run the domain size calculations identically on the simulations as for the experimental data, we up-sampled the cell type classification grid for each simulation to the experimental spatial resolution using the imresize() function.

### Experimental cell sorting assays

#### Cell lines and culture

Cell lines were cultured at 37 °C with 5% CO_2_. HEK293T (ATCC; CRL-3216) cells were cultured in DMEM (ATCC) supplemented with 9% fetal bovine serum (FBS; ATCC; 30-2020), Penicillin-Streptomycin (Pen-Strep; ATCC; 30-2300) at a final concentration of 90 I.U./mL penicillin and 90 µg/mL streptomycin, and 1.8 mM L-Glutamine. The FBS solution was heat-inactivated before use by incubating it at 56 °C in a water bath for 30 minutes prior to addition to the media. L929 (ATCC; CCL-1) cells were cultured in EMEM (ATCC; 30-2003) supplemented with 9% horse serum (Thermo Fisher Scientific; 16050122) and Pen-Strep (ATCC; 30-2300) at a final concentration of 90 I.U./mL penicillin and 90 µg/mL streptomycin. For routine passaging of both cell lines, cells were washed with Dulbecco’s Phosphate Buffered Saline (D-PBS; ATCC; 30-2200), incubated with 20 µL/cm^2^ Trypsin-EDTA solution (ATCC; 30-2101) for 5 minutes at 37 °C, and resuspended by addition of 4-fold excess (volume-wise) complete media. Cells were passaged at a ratio of 1:10 every 3–4 days, using T-75 flasks (Corning; 353136) for both HEK293T and unmodified L929 lines, and six-well plates (Corning; 3516) for Cdh-expressing L929 lines, whose generation is described below.

#### Generation of Cdh-P2A-FP–containing lentiviral transfer vectors

All cloning steps were performed by Gibson Assembly using the NEBuilder® HiFi DNA Assembly Master Mix (NEB; E2621L). First, a stop codon (TAA) was added downstream of the EGFP ORF in the pLJM1–EGFP lentiviral transfer vector (Addgene; plasmid #19319). This resultant plasmid (pLJM-EGFP_v2) was then used to generate a pLJM1-mRFP1 plasmid by replacing the EGFP coding sequence (CDS) with the mRFP1 CDS, obtained by PCR-amplification from the pcDNA-mRFP1 plasmid (Addgene; plasmid #13032). Finally, each of pLJM1-mRFP1 and pLJM1-EGFP_v2 plasmids was used to generate the pLJM1-Cdh1-P2A-EGFP and the pLJM1-Cdh3-P2A-mRFP1 plasmids by replacing the start codon of each of the fluorescent proteins with the stop codon– removed CDS of either mouse *Cdh1* or *Cdh2,* followed by the P2A self-cleaving peptide sequence (N-GSGATNFSLLKQAGDVEENPGPGSG-C). The *Cdh1* CDS and the *Cdh2* CDS were each PCR-amplified from the pBATEM2 (BCCM consortium; LMBP 3585) and pBact-Pcad (BCCM consortium; LMBP 2766) plasmids, respectively.

#### Generation of Cdh-expressing L929 cell lines

*<u>Cdh1–P2A–mRFP1</u>*<u>- and *Cdh3–P2A–EGFP*</u> - expressing L929 cell lines were generated by lentiviral transduction, using a protocol adapted from (Kleaveland et al., 2018[https://doi.org/10.1016/j.cell.2018.05.022]). Lentiviral particles were produced by reverse-transfection of HEK293T cells, which was performed as follows: 0.938 µg of the psPAX2 packaging vector (Addgene; plasmid #12260), 0.469 µg of the pMD2.G envelope vector (Addgene; plasmid #12259), and 1.41 µg of one of the two a pLJM1-derived transfer vector described in the previous section were mixed with 5.5 µL of P3000 (L3000015) in 50 µL of Opti-MEM (Thermo Fisher Scientific; 31985062), and then combined with 8.25 mL of Lipofectamine 3000 (Thermo Fisher Scientific; L3000015) in 50 µL of Opti-MEM, according to the manufacturer’s instructions. Each plasmid–Lipofectamine 3000 solution was then added to a separate well within a 6-well plate, after which 1.5 million HEK29T cells in 2 mL of complete media were added to each well and the plate was incubated at 37 °C and 5% CO_2_ for 48 hours.

Viral particles were collected by removing the media from the transfected HEK293T cells, and twice centrifuging at 500 G for 10 min at 25 °C to pellet debris. Infection of L929 cells was performed by adding 250 µL of the cleared viral supernatant to L929 cells that had been plated 24 h earlier in a 12-well plate at a density of 1 million cells in 750 µL of complete media per well, followed by addition of polybrene (Millipore Sigma; TR-1003-G) to a final concentration of 8 µg/mL. Plates were then centrifuged at 1,200 G for 90 minutes at 25 °C, and then incubated at 37 °C and 5% CO_2_ for 48 hours.

Stably transduced L929 cells were then selected by culturing the cells for 1 week with complete media containing 4.3 µg/mL Puromycin (Thermo Fisher Scientific; A1113803). Low- and high-expression populations were sorted from each of the two stable lines using the BD FACSAria II flow cytometer (BD Biosciences), selecting the 5–20^th^ percentile of the fluorescence distribution to be the low-expression population, and the 80–95^th^ percentile to be the high-expression population.

Note: Given that (1) the *Cdh2*- and *Cdh3*-expressing cell lines exhibited overall similar levels of fluorescence (**Fig. 6 – Supp.** Fig. 1) and therefore also presumably expressed similar levels of cadherin, and (2) *Cdh2* and *Cdh3* have comparable homophilic binding affinities and comparable heterophilic binding affinities to *Cdh1* in vitro(*107*), it was perhaps surprising that conditions in which *Cdh3*- and *Cdh1*-expressing cells were mixed sorted consistently better than conditions in which *Cdh2*- and *Cdh1*-expressing cells were mixed (**Fig. 6a**).These results suggest that *Cdh3* and *Cdh2* may be regulated differently in L929 cells in terms of their turnover rates, clustering dynamics, or their anchoring to the actin cytoskeleton(*108*). Recent work in cells has shown that the binding affinity of the extracellular domains of cell adhesion molecules is less important for defining the adhesive specificity between cells than is the intracellular domain(*108*).

#### Cell-sorting assay

Cell-sorting assays were performed by first mixing either Cdh1-P2A-mRFP1^high^ L929 cells with Cdh2-P2A-EGFP^high^ cells or Cdh1-P2A-mRFP1^low^ L929 cells with Cdh2-P2A-EGFP^low^ cells at a 1:1 ratio, and adding 80,000 cells of this mixture to a well of a 384-well round-bottom ultra-low-attachment spheroid microplate (Corning; 4516) in 100 µL of complete media. After loading all samples into the microplate, the plate was centrifuged at 150 G for 10 minutes at 25 °C. The positions of the wells used within the microplate chosen to be located as close to the center of the microplate as possible, so that this centrifuging step would pellet the cells close to the center of each of the round-bottom wells. After pelleting, the cells were maintained at 37 °C and 5% CO_2_ using a microscope stage–top incubator (Tokai Hit).

#### Microscopy

Time-lapse images of each assay were acquired using an inverted Nikon A1R point-scanning confocal microscope and NIS-Elements AR 4.51 (Nikon) software, concurrently illuminating the samples with 488 nm and 561 nm lasers to image the EGFP and mRFP1 signal, respectively. Emitted light was collected with a 10× Plan Apo DIC N1 objective (Nikon; numerical aperture = 0.45; working distance = 4000 µm) and two photomultiplier (PMT) detectors at 1024×1024 *xy* pixels. Volumetric images 1270×1270×200 µm (*xyz*) in size with 1.24×1.24×10 µm/pixel (xyz) spatial resolution were collected every 30 minutes for 24 hours.

### Pre-processing of the experimental data for domain size calculations

All image analysis was performed in MATLAB.

*Max intensity z-projections (MIP) –* Raw 3D+time images were loaded using bioformats tools, and maximum intensity Z-projections on the were performed on each 3D image volume using the max() function.

#### Cell type classification

The raw max intensity z-projections were converted to double precision using MATLAB’s im2double() function. The channels were then histogram matched to enforce comparable distributions of pixel intensity values using the imhistmach() function, so that one channel would not be favored over the other during cell type classification. The images were then segmented by thresholding using a user-defined threshold that was specific to each dataset (typically between 0.002 and 0.0035) using the imbinarize() function. Holes in the segmented images that were smaller than one cell area (40 pixels) were filled using the bwareaopen() function. After this, each pixel was classified. Pixels that were segmented in only one channel were assigned the cell type representing that channel (i.e., given a value of either 1 or 2, representing the EGFP or mRFP1 channel, respectively). For pixels that were segmented in both channels, the pixel was assigned to the channel with the higher histogram-matched pixel intensity. Pixels that were not segmented in either channel were given an arbitrary alternative value of 1.5. This process resulted in a classified image the same size as the original image, where each pixel was assigned a value of 1 (cell type #1), 2 (cell type #2), or 1.5 (no cell).

### Experimental fluorescence intensity calculations

For fluorescence intensity calculations (**Fig. 6 - Supp.** Fig. 1), the median pixel value was averaged across all pixels in the raw MIP image for the first timepoint and calculated for each channel using the median() function.

### Sorting quantification

#### (Simulations only) Percent same cell type neighbor calculations

Neighbors were defined as any two cells sharing faces (but not corners, i.e., 4-pixel connectivity). For each cell, the number of neighbors of the same cell type was counted, divided by the total number of neighbors, and then multiplied by 100 in order to calculate the percent of same-cell-type neighbors. This percent of same cell type neighbors was than averaged across all cells in the tissue to calculate the degree of sorting.

#### Domain size calculations (overview)

Domain size calculations were performed by image skeletonization, a common procedure that finds the “backbones” that run along the centers of connected domains within a segmented image. Once the skeleton of the cell type domains was extracted, the domain size was calculated as twice the closest distance from each point along the skeleton to the edge of the domain (**Fig. 1 – Supp.** Fig. 1). Domain size calculations were performed separately for each cell type.

#### Domain size calculations (detailed)

For each cell type, a binary image was generated that was given a value of 1 for every pixel classified as that cell type, and all other pixels were set to 0 (**Fig. 1 – Supp.** Fig. 1a). In this binary image, holes smaller than the average cell area were filled using the bwareaopen() function, and closed objects smaller than the average cell area were removed using the bwareafilt() function (**Fig. 1 – Supp.** Fig. 1b). The boundaries of each object in the binary image were then smoothed over a distance of one cell diameter using the conv2 function and then thresholded to create a binary image again (**Fig. 1 – Supp.** Fig. 1b).

From this processed binary image, the skeleton along each cell type domain was extracted using the bwskel() function, and the distance of each point along the skeleton to the nearest edge of the domain was calculated using the bwdist() function. Wide domains contain more pixels than narrow domains, but are counted equally by skeleton-based calculations. To ensure each pixel in the image was represented equally, we then weighted the domain size at each point along the skeleton by the number of pixels in that domain.

The analyses described so far worked well for relatively elongated, thin domains, but failed for thick domains with many small holes (small being relative to the domain size). To remove holes smaller than the domain size, the entire process was repeated, this time filling holes and smoothing each object not by the average cell diameter, but by the average domain size of that binary object.

Finally, the analyses up to this point failed for domains with circular or square-shaped domains, as these objects do not have well-defined skeletons. Therefore, a final check was performed on each object, such that for objects with a minor axis length/major axis length > 0.85 and a solidity > 0.9, the domain size was instead calculated as the major axis length and weighted by the total number of pixels in that object.

### Neighbor exchange rate calculations

For the neighbor exchange rate calculations (**Fig.1f**) – first, a list of the identities of each neighbor for each cell for each timepoint was generated. Neighbors were defined as all cells within one cell diameter of that cell (all neighbors that share a face, but not those that share corners). Once the neighbors were identified, the algorithm counted (for each cell) how many old neighbors were lost, and how many new neighbors were gained between timepoints. The total number of neighbors exchanged in each timepoint was equal to the sum of the number of neighbors lost plus the number of neighbors gained.

To calculate a neighbor exchange rate, a cumulative sum was performed on the number of neighbors exchanged over time, in order to calculate the cumulative number of neighbors exchanged since the first timepoint (a much smoother metric than raw number of neighbors exchanged per timepoint). The cumulative number of neighbors exchanged was then averaged across all cells to calculate the mean cumulative number of neighbors exchanged. Finally, a moving linear regression (using a 15-timepoint window) was performed on the mean number of cumulative number of neighbors exchanged vs time curve. From this moving linear regression, the instantaneous neighbor exchange rate within the tissue over time (**Fig. 1f**) was determined as the instantaneous slope of the curve at each timepoint.

### Fitting of the simulations to the experimental data

#### Parameter scan of simulations

As discussed earlier in the text, because tissue fluidity in the model is determined by two major ratios—the diffusion limited neighbor exchange rate (***k_0_ = E_M_ / 3πηd^3^***) and the exponent (***E_M_ / E_A_***)—different parameter choices with identical ratios will yield an identical sorting behavior. Therefore, we performed a parameter scan on these two ratios in the model.

As a simplifying assumption to limit the number of parameters being fit to the model, the heterotypic adhesion (***E_het_***) was assumed to be zero, and the homotypic adhesion (***E_homo_***) was assumed to be equal between the two cell types being mixed, and the motility energy (***E_M_***) was assumed to be equal between the two cell types being mixed. As a consequence of the heterotypic adhesion (***E_het_***) being zero, the total adhesion energy (***E_A_***) is entirely determined by the homotypic adhesion energy (***E_homo_***).

This leaves only three major parameters that could be varied: The homotypic adhesion energy (***E_homo_***), the motility energy (***E_M_***), and the viscosity (***η***). Because different parameter choices with identical ratios will yield an identical sorting behavior, we therefore varied the two simplified ratios: The ratio of the motility energy to the homotypic adhesion energy (***E_M_ / E_homo_***), and the ratio of the homotypic adhesion energy to the viscosity (***E_homo_ / η***).

The unitless ratio of the exponent (***E_M_ / E_homo_***) varied between 0.25 and 1.5 in 0.05 intervals. The ratio of the homotypic adhesion energy to the viscosity (***E_homo_ / η***) varied from 10^3^ - 10^7^ ***k_B_T*** / Pas. Assuming a viscosity of ***η*** = 10 Pa s, this means that the homotypic energy ranged from 10^4^ -10^8^ ***k_B_T***.

For each set of parameters, simulations were run for 48 hours, and the mean and standard deviation of domain size was calculated for each timepoint in the simulations. 5 simulation replicates were performed for each choice of parameters

#### Magnitude-weighted least absolute deviations (WLAD) fitting of simulations to the experimental data –

Each experimental dataset was compared to all simulations run in the parameter scan. Because the experiments exhibited a lag phase in which very little sorting (as measured by the domain size) occurred during the first 2 hours, hours 0-18 of the simulations were fit to hours 2-20 of the experiments. The fitting algorithm obtained the best fit simulation parameters based on the magnitude-weighted least absolute deviations (LAD) between the domain size curves for the simulations and the experiments. LAD was used instead of least squares (LS) because LAD is less sensitive to outliers and yielded more consistent fit results between the experimental replicates than LS, though LS fitting yielded similar results. First, the absolute deviation, ***AD(t)***, between the mean domain size, ***MEAN(t)***, and the standard deviation in domain size, ***STD(t)***, for each simulation, ***MEAN_Sim_(t)*** and ***STD_Sim_(t)***, and each experiment, ***MEAN_Exp_(t)*** and ***STD_Exp_(t)***, was calculated as ***AD_MEAN_(t) = |MEAN_sim_(t)-MEAN_Exp_(t)|*** and ***AD_SD_(t) = |SD_sim_(t)-SD_Exp_(t)|*** for each timepoint. To account for the fact that simulations with larger domain sizes will naturally exhibit larger fluctuations in the domain size, the absolute deviations were then weighted by the inverse of (i.e., normalized to) the domain size in the simulations at each timepoint, or ***WAD_MEAN_(t) = AD_MEAN_(t)/MEAN_sim_(t)*** and ***WAD_SD_(t) = AD_SD_(t)/SD_sim_(t)***. The weighted absolute deviations were then averaged across simulation replicates, or ***<WAD(t)>_reps_***, and then summed across time ***WADT_l_*** = ∑***<WAD(t)>_reps_***. The final cost function was defined as the sum of the WAD for the mean and standard deviation in the domain size, ***C = WADT_Mean_ + WADT_SD_***. The best fit simulation parameters were defined as those with the lowest cost value. Heatmaps of the cost function show that the best fit parameters occupied reasonably well-defined minima in parameter space (**Fig. 6 – Supp.** Fig. 3). The clustering of the best fit parameters for the experimental replicates also suggests that these values are fairly well-constrained (**Fig. 6d**).

*Note:* While the parameter scan and fitting algorithm in its current form provides good quantitative fits to the experimental data (Fig. 6 – Supp Fig. 2), it is important to note that the assumption of the heterotypic binding energies being zero and the homotypic adhesion being equal between the two cell types is not strictly consistent with the literature. In fact, it has been shown in vitro (with supporting evidence from cell sorting assays) that the extracellular binding domains of *Cdh1* bind more strongly in heterotypic interactions with either Cdh2 or Cdh3 than in homophilic interactions with itself, such that the binding interaction strengths are ordered as [*Cdh2-Cdh2* and *Cdh3-Cdh3*] > [*Cdh2-Cdh1* and *Cdh3-Cdh1*] > [*Cdh1-Cdh1*] (*107*). Future iterations using more information about the sorting pattern than just the domain size (e.g., relative domain sizes between the cell types, the overall spatial localization of different cell types within the tissue) could be adapted to more fully estimate the differential homotypic and heterotypic binding affinities between the cell types.

## Materials availability

All cell lines and plasmids will be made available upon reasonable request.

## Data and code availability

All raw imaging data (∼ 300 GB of images stored as Nikon ‘.nd2’ files), processed imaging data (∼ 10 GB of maximum intensity projected movies and segmented movies stored as ‘tiff’ files, and calculated domain sizes stored as ‘.mat’ files), and simulation data (stored as MATLAB ‘.mat’ files) will be made available upon reasonable request. All simulation and data analysis code (stored as MATLAB ‘.mat’ files) are freely available on Gitlab: <https://github.com/megasonlab/Garner_etal_2025_Fluidity_Sorting>

## Acknowledgements

We thank Lisa Manning, Alon Oyler-Yaniv, and the members of the labs of Sean Megason, Allon Klein, Olivier Pourquié, and Marcos Simoes-Costa for helpful discussions and feedback.

## Author Contributions

Conceptualization, R.M.G.; Methodology, R.M.G. and S.E.M.; Software, R.M.G.; Validation, R.M.G. and S.E.M.; Formal Analysis, R.M.G.; Investigation, R.M.G. and S.E.M.; Resources, R.M.G. and S.E.M.; Data curation, R.M.G. and S.E.M.; Writing – Original Draft, R.M.G..; Writing – Review & Editing, R.M.G., S.G.M., S.E.M., A.M.K.; Visualization, R.M.G.; Supervision, S.G.M., A.M.K.; Project Administration, R.M.G., S.G.M., S.E.M., A.M.K.; Funding Acquisition, S.G.M., A.M.K.;

## Declaration of Interests

A.M.K. is a cofounder of Somite Therapeutics, Ltd.

## Declaration of generative AI and AI-assisted technologies

The authors declare no use of generative AI and AI-assisted technologies.

## Funding

This work was supported by grants to S.G.M. (NIH R01GM107733, NIH R01DC015478, NIH R01GM154339), a grant to S.G.M. and A.M.K. (NIH R01EY036381), Helen Hay Whitney Fellowships to R.M.G. and S.E.M., and a HHMI Hanna Gray Fellow Finalist gift to R.M.G.

---

**Figure 1 – Supplemental Figure 1.**
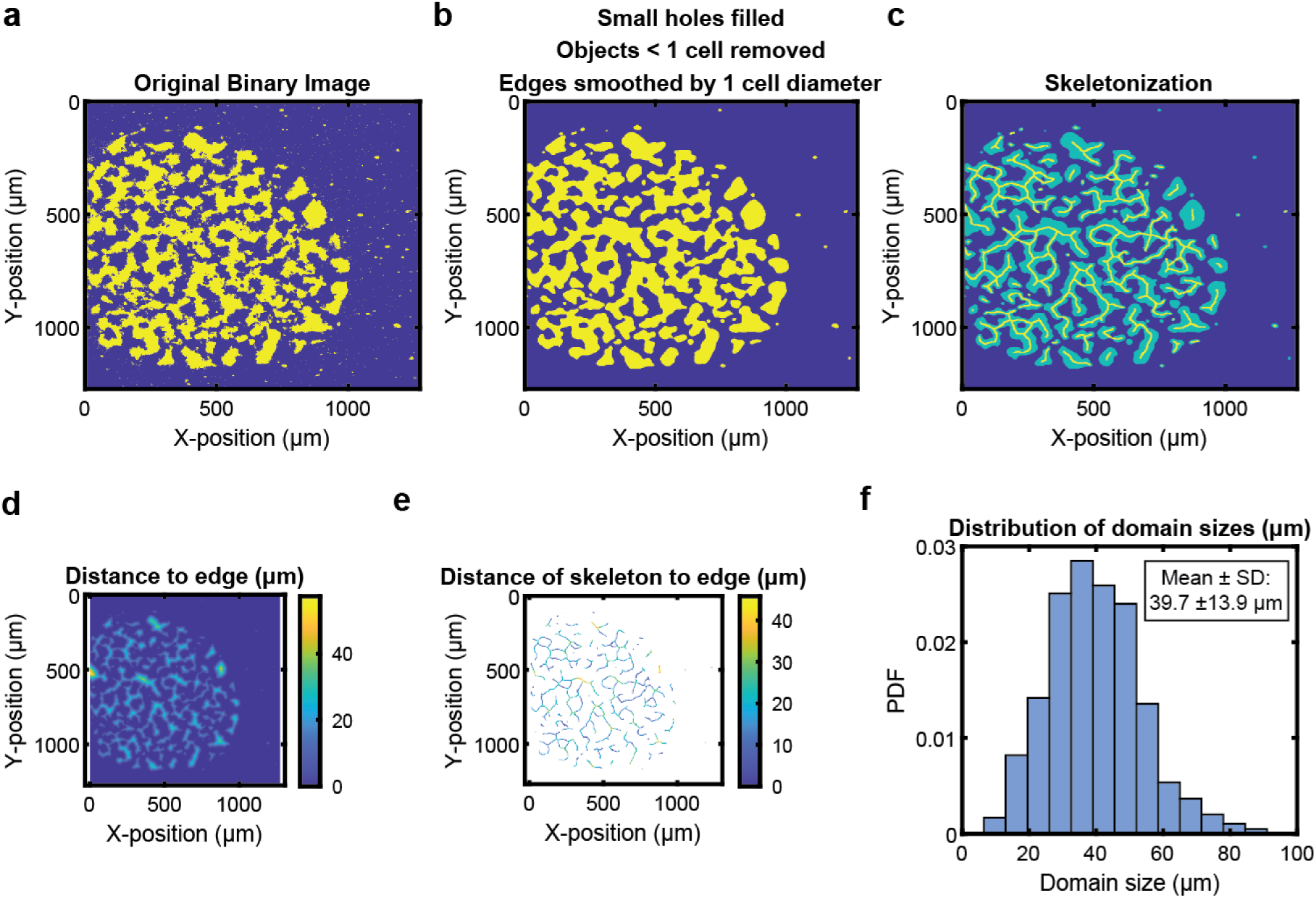
Domain size calculations. (**a-f**) Overview of the steps of the domain size calculation. See ***Methods*** for a detailed description of each panel.

**Figure 1 – Supplemental Figure 2.**
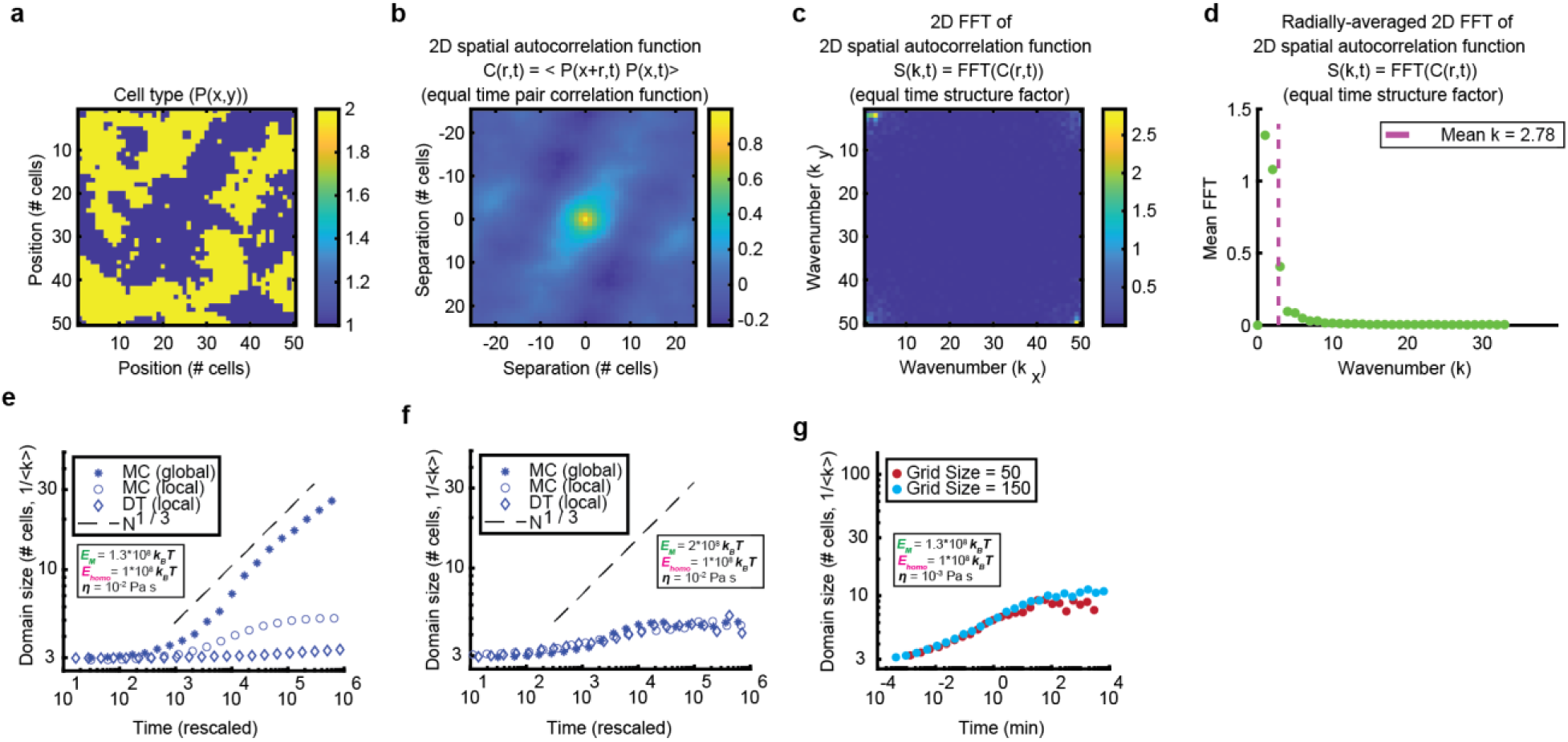
Domain size calculations. (**a-d**) Overview of the steps of the domain size calculations for the purpose of comparing simulation methods. From the cell type matrix (a), the 2D spatial autocorrelation was computed (b), and then the Fourier transform of the autocorrelation function was determined to calculate the structure factor (c), which was then radially averaged (d). The domain size was then calculated as the mean structure factor weighted wavenumber. (e) Comparison of the three simulation methods, showing that simulations allowing only direct (i.e., sharing faces) neighbor exchange (MC Local and DT local) exhibit sub-t^1/3^ scaling, capturing the short-time glassy behavior of the system. (f) Comparison of the three simulation methods in the high motility regime, showing that all simulation methods converge and saturate at domain sizes much smaller than the grid size. (g) Comparison of simulations with a grid size of 50 and 150 cells, showing that growth saturation at domain sizes much smaller than the grid size is not due to finite size effects.

**Figure 2 - Supplemental Figure 1.**
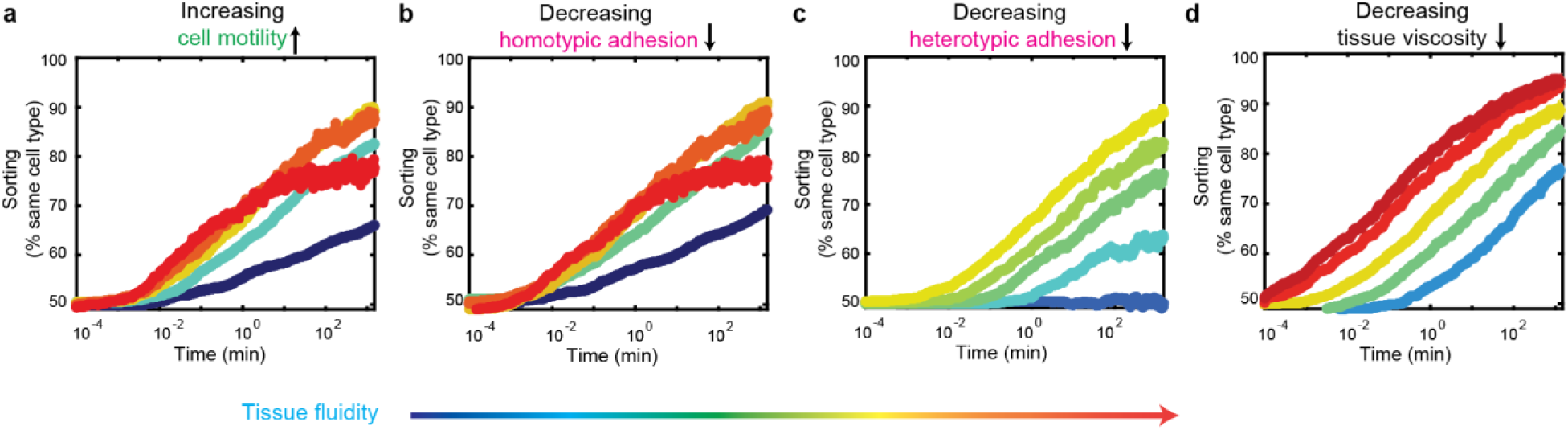
The sorting rate increases with increasing tissue fluidity. (a-d) Degree of sorting (i.e., average percent of a cell’s neighbors that are of the same cell type) plotted as a function of time for the exact same simulations presented in Fig. 2m-p, with increasing tissue fluidity from blue to red. Tissue fluidity was increased either by (a) increasing motility energy, (b) decreasing homotypic adhesion, (c) decreasing heterotypic adhesion, or (d) decreasing viscosity. The colormap is identical to that in Fig. 2. In all cases the rate of sorting increases (i.e., the steepness of the curve increases and/or the curve shifts to the left, such that curves with higher tissue fluidity reach the same degree of sorting at earlier timepoints than curves with lower tissue fluidity).

**Figure 3 – Supplemental Figure 1.**
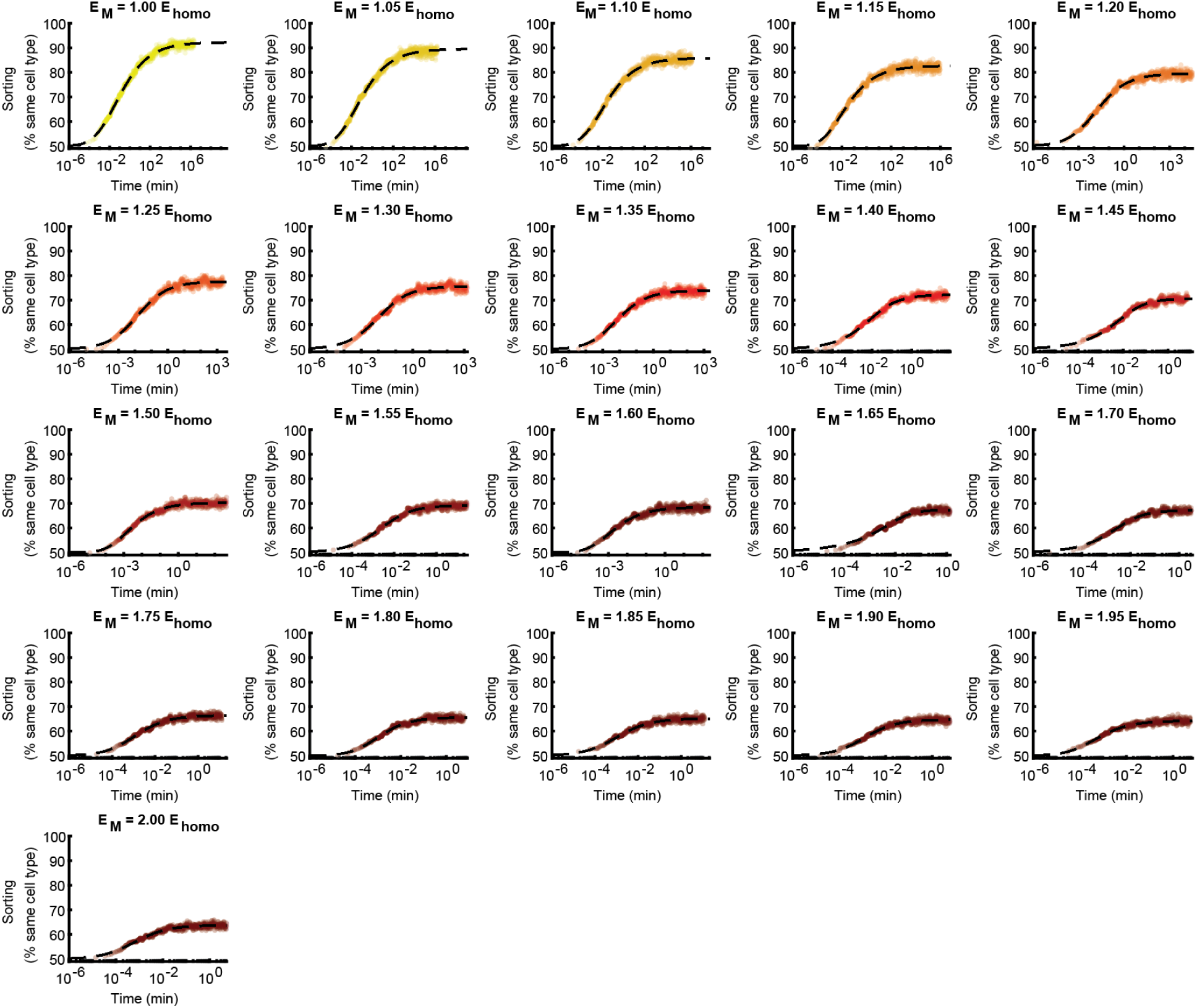
Best fit curve for each simulation. (**a**) Simulations shown in in Fig. 3b**-e** were allowed to run until they reached steady state. At this point, the sorting vs time curve (colored circles) was fit to an asymmetric Hill function (black dotted line). The color of the curve indicates the choice of motility energy, using an identical colormap as in Fig. 3b**-e**. Note the differences in scale along the x-axis.

**Figure 6 – Supplemental Figure 1.**
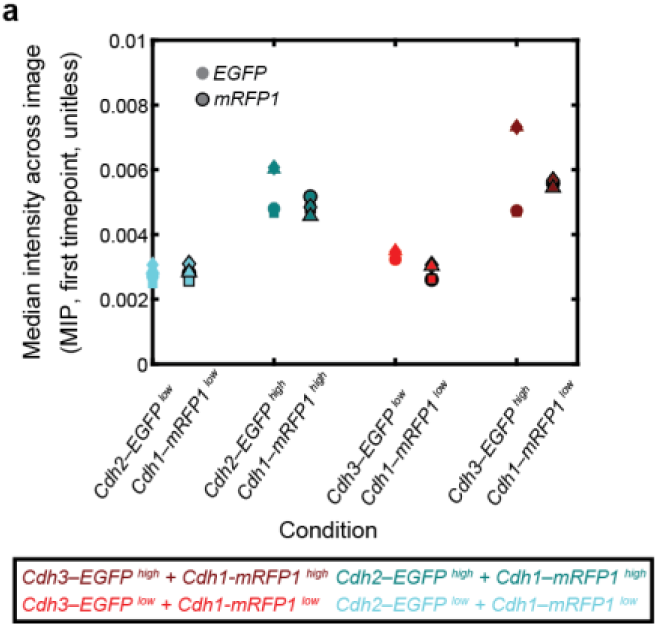
Median fluorescence intensity for each cell type in each experimental condition. (**a**) L929 cells co-expressing either high or low levels of *Cdh2* (N-cadherin) or *Cdh3* (P-cadherin) and *EGFP* (green) were mixed in equal proportions with cells co-expressing either high or low levels of *Cdh1* (E-cadherin) and *mRFP1* (magenta) and imaged by confocal time-lapse microscopy. From the maximum intensity projection of the first timepoint for each condition, the median fluorescence intensity across the entire image was calculated for both the *EGFP* (open circles) and *mRFP1* (closed circles) channels. The color indicates the experimental condition, and the two cell populations mixed in a single condition are grouped together along the x-axis. The x-label indicates which cell population corresponds to each channel within a given condition. The markers indicate different experimental replicates of a given condition. There were four replicates in each condition.

**Figure 6 – Supplemental Figure 2.**
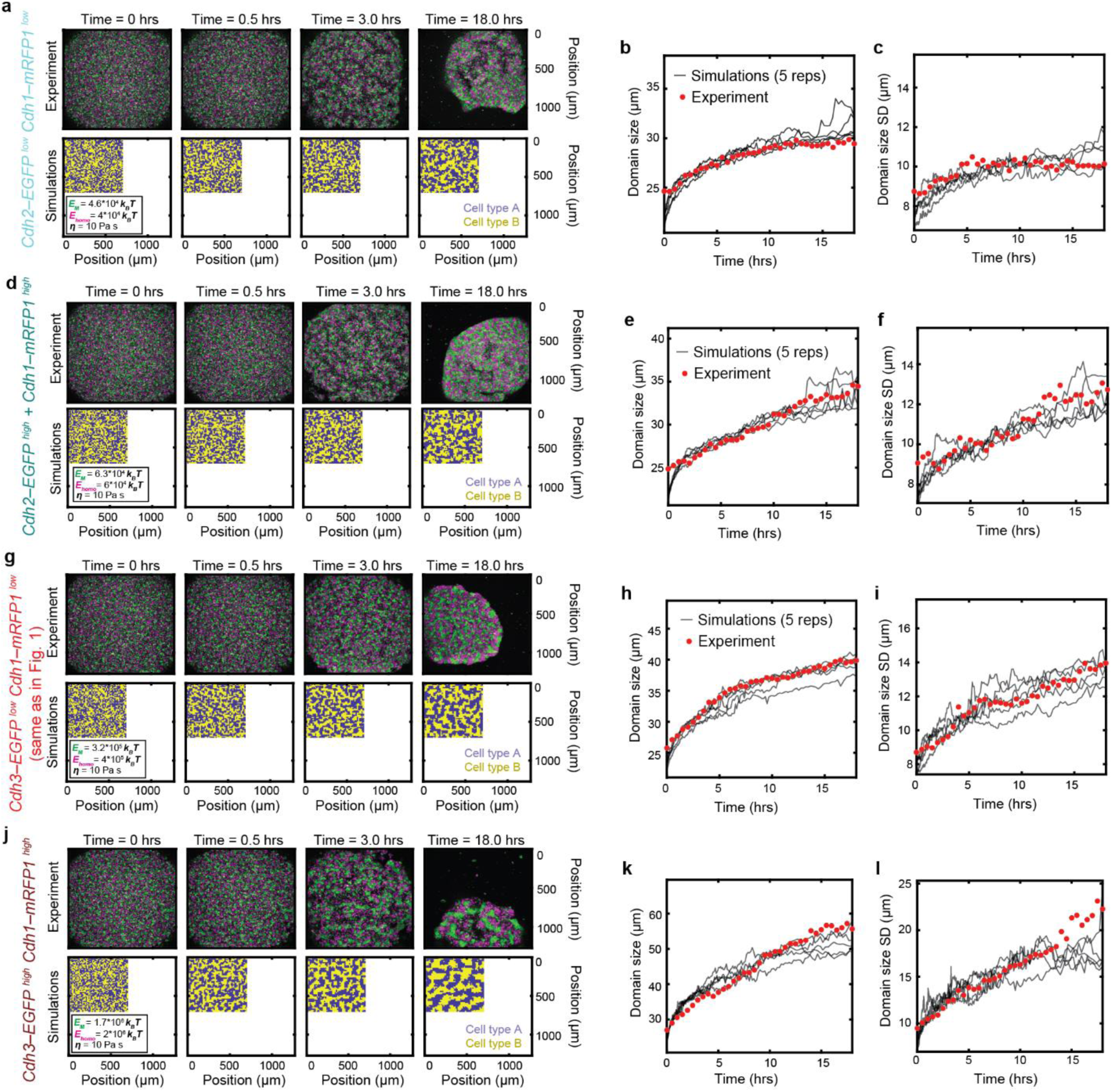
Examples of best fit for each experimental condition. (**a,d,h,j**) Time-lapse montage of the experimental cell-sorting assay (*top*) and its associated best-fit simulation (*bottom*). *Top:* L929 cells co-expressing either high or low levels of *Cdh2* (N-cadherin) or *Cdh3* (P-cadherin) and *EGFP* (green) were mixed in equal proportions with cells co- expressing either high or low levels of *Cdh1* (E-cadherin) and *mRFP1* (magenta) and imaged by confocal time-lapse microscopy. Images represent maximum intensity projections. *Bottom:* Best-fit simulations with displayed as a heat map of cell type. Best fit parameters are listed in the inset legend. (**b,c,e,f,h,i,k,l**) Mean (**b,e,h,k**) and standard deviation (**c,f,i,l**) of the same-cell-type domain size over time for the experiments (red dots) and for 5 replicate best-fit simulations (grey lines). (**a-c**) *Cdh2–EGFP*^low^ cells mixed with *Cdh1*–*mRFP1*^low^ cells, (**d-f**) *Cdh2–EGFP*^high^ cells mixed with *Cdh1–mRFP1*^high^ cells, (**g-i**) *Cdh3*–*EGFP^low^* cells mixed with *Cdh1*–*mRFP1^low^*cells (**j-l**) *Cdh3–EGFP*^high^ cells mixed with *Cdh1*-*mRFP1^high^*cells,

**Figure 6 – Supplemental Figure 3.**
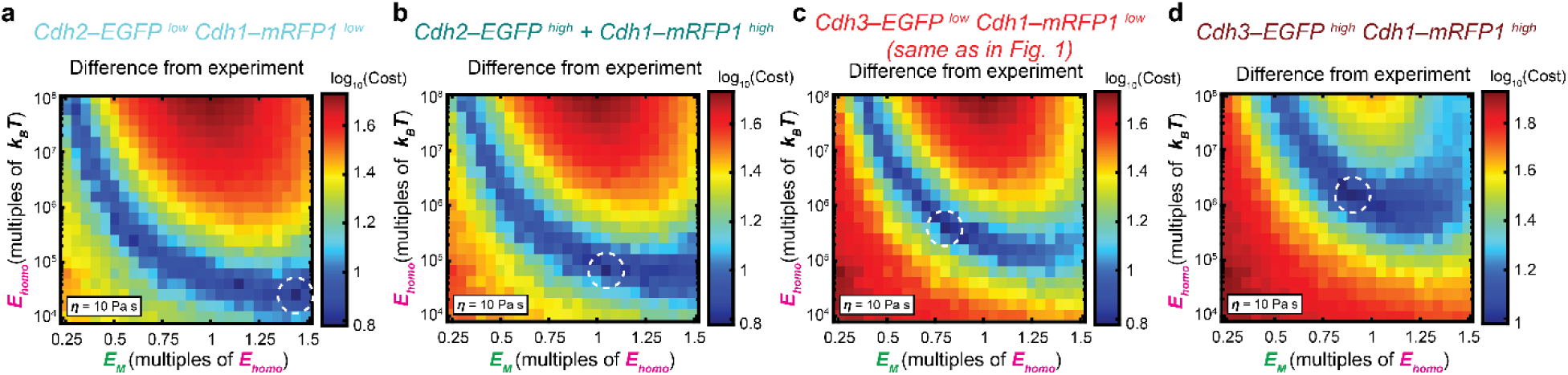
Cost values for fitting the simulations to the experimental data. (**a-d**) Heatmap of the magnitude-weighted least absolute differences between the experiment and all possible parameter choices for the simulations, averaged over 5 simulation replicates. The best fit simulation parameters (i.e., minima) are denoted by dashed white circles. (**a**) *Cdh2–EGFP*^low^ cells mixed with *Cdh1*–*mRFP1*^low^ cells, (**b**) *Cdh2–EGFP*^high^ cells mixed with *Cdh1–mRFP1*^high^ cells, (**c**) *Cdh3*– *EGFP^low^* cells mixed with *Cdh1*–*mRFP1^low^* cells, (**d**) *Cdh3–EGFP*^high^ cells mixed with *Cdh1*-*mRFP1^high^*cells

**Figure 6 – Supplemental Figure 4.**
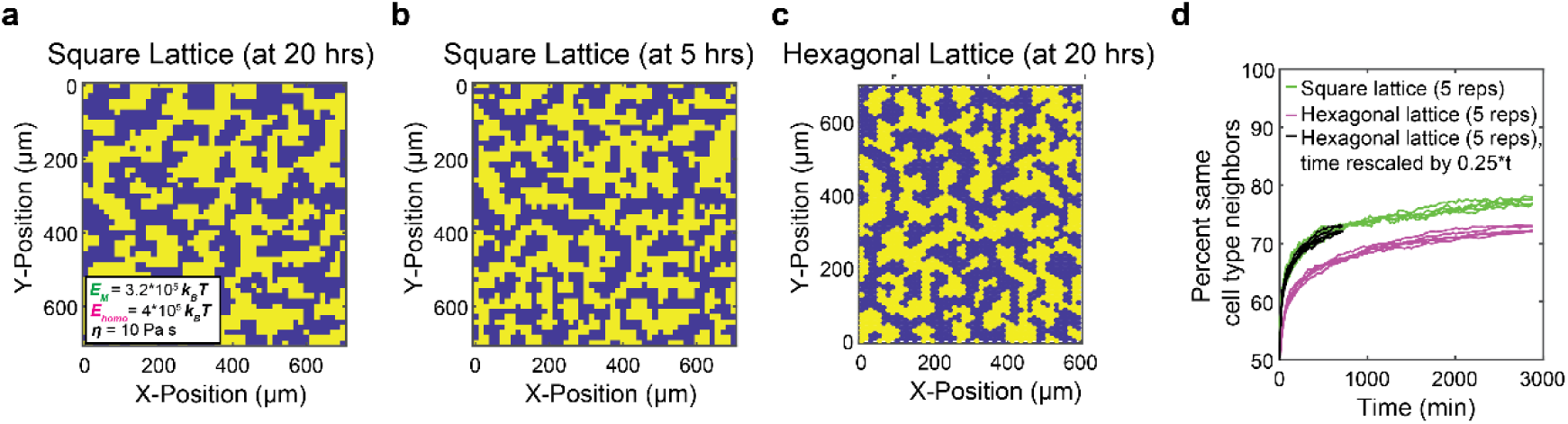
Lattice geometry rescales sorting time. (**a-d**) Comparison of a square and hexagonal lattice for simulations with the same parameters used in Fig. 1g and Fig. 6c (second from right). (**a**) Heatmap of the cell type arrangement on a square lattice after 20 hours of sorting time. (**b**) Heatmap of the cell type arrangement on a square lattice after 5 hours of sorting time, best mapping to the degree of sorting on a hexagonal lattice shown in (c). (**c**) Heatmap of the cell type arrangement on a hexagonal lattice after 20 hours of sorting time. (**d**) Degree of sorting plotted as a function of time for a square lattice (*green*), hexagonal lattice (*pink*), and a hexagonal lattice with time rescaled to 0.25\****t*** (*black*), showing close overlap with the square lattice.

**Table S1.** Derived parameters. Parameters listed are the default used for the simulations.

| Notation | Meaning | Default Value | Typical Range | Source |
| --- | --- | --- | --- | --- |
| $k_B$ | Boltzmann constant | 0.0138 pN nm K<sup>-1</sup> | – | – |
| $T$ | Temperature | 310.15 K (37 °C) | 25-40 °C | – |
| $r$ | Cell radius | 7,000 nm | – | – |
| $d$ | Distance between cell centers | $2*r = 14,000$ nm (confluent) | – | – |
| $\eta_w$ | Viscosity of water | $10^{-6}$ pN ms nm<sup>-2</sup> (10<sup>-3</sup> Pa s) | – | – |
| $\eta$ | Effective Viscosity | $10^{-2}$ pN ms nm<sup>-2</sup> (10 Pa s) | $10^{-6}$-$10^0$ pN ms nm<sup>-2</sup> (10<sup>-3</sup>-10<sup>3</sup> Pa s) | (25) |
| $E_{A, homo}$ | Homotypic adhesion energy (per cell pair) | $10^8 k_B T$ | $0$-$10^8 k_B T$ | |
| $E_{A, het}$ | Heterotypic adhesion energy (per cell pair) | 0 | $0$-$10^8 k_B T$ | |
| $E_M$ | Effective migratory temperature of cell | $10^8 k_B T$ | pN nm | |
| $N_{homo}$ | Number of homotypic neighbors for a given cell | - | 0-4 neighbors | – |
| $N_{het}$ | Number of heterotypic neighbors for a given cell | – | 0-4 neighbors | – |

| Notation | Meaning | Value | Units | Source |
| --- | --- | --- | --- | --- |
| $k_B T$ | Thermal energy | $k_B T$ | pN nm | – |
| $\gamma$ | Viscous drag on a cell | $6\pi\eta R$ | pN ms nm<sup>-1</sup> | Stokes' drag |
| $E_T$ | Total adhesion energy for a given cell | $N_{homo}E_{homo} + N_{het}E_{het}$ | pN nm | |
| $E_A$ | Total adhesion energy for a given neighbor pair | $E_T^1 + E_T^2$ | | |
| $k_0$ | Neighbor exchange rate (for each individual neighbor pair) in the absence of adhesion | $E_M / \gamma d^2$ | ms<sup>-1</sup> | Diffusion-limited |
| $k$ | Neighbor exchange rate (for each individual neighbor pair) | $k_0 \exp(-E_A/E_M)$ | ms<sup>-1</sup> | Diffusion-limited |

## Notes

### Summary of Updates

This version of the manuscript has been revised in response to reviewers, and has been accepted for publication at Biophysical Journal. The revised manuscript includes substantial revisions and clarifications throughout the main text, over a dozen additional citations, and two additional supplementary figures.

